# A role of Lck annular lipids in the steady upkeep of active Lck in T cells

**DOI:** 10.1101/2022.03.18.484902

**Authors:** Nicla Porciello, Deborah Cipria, Giulia Masi, Anna-Lisa Lanz, Edoardo Milanetti, Alessandro Grottesi, Duncan Howie, Steve P. Cobbold, Lothar Schermelleh, Hai-Tao He, Marco D’Abramo, Nicolas Destainville, Oreste Acuto, Konstantina Nika

**Affiliations:** T Cell Signalling Laboratory, Sir William Dunn School of Pathology. Oxford University, Oxford, OX2 3RE, United Kingdom; Department of Physics, University of Rome “La Sapienza”, 00185, Rome, Italy; CINECA - Italian Computing Centre (ICC). 00185 Rome, Italy; Sir William Dunn School of Pathology. Oxford University, Oxford, OX2 3RE, United Kingdom; Micron Advanced Bioimaging Unit; Department of Biochemistry, Oxford University, OX1 3QU, United Kingdom; Centre d’Immunologie de Marseille-Luminy, Aix-Marseille Université, Marseille, France; Department of Chemistry, University of Rome “La Sapienza”, 00185, Rome, Italy; Laboratoire de Physique Théorique, Université de Toulouse, CNRS, UPS, France; Department of Biochemistry, School of Medicine. University of Patras, Greece

**Keywords:** Lck, CD45, membrane anchor, annular lipids, membrane lateral organisation

## Abstract

Theoretical work suggests that collective spatiotemporal behaviour of integral membrane proteins (IMPs) can be modulated by annular lipids sheathing their hydrophobic moiety. Here, we present evidence for this prediction in a natural membrane by investigating the mechanism that maintains steady amount of active isoform of Lck kinase (Lck_A_) by Lck trans-autophosphorylation offset by the phosphatase CD45. We gauged experimental suitability by quantitation of CD45 and Lck_A_ subcellular localisation, Lck_A_ generation as a function of Lck and pharmacological perturbation. Steady Lck_A_ was challenged by swapping Lck membrane anchor with structurally divergent ones expected to substantially modify Lck annular lipids, such as that of Src or the transmembrane domains of LAT, CD4, palmitoylation-defective CD4 and CD45, respectively. The data showed only small alteration of Lck_A_, except for CD45 hydrophobic anchor that thwarted Lck_A_, due to excessive lateral proximity to CD45. The data are best explained by annular lipids facilitating or penalising IMPs’ lateral proximity, hence modulating IMPs protein-protein functional interactions. Our findings can contribute to improve the understanding of biomembranes’ organisation.

## Introduction

Cell decisions in response to environmental cues initiate by events choreographed at the plasma membrane (PM) by integral membrane proteins (IMPs). IMPs’ function is influenced by membrane lipids and vice-versa, producing a highly dynamic and complex two-dimensional milieu. Embedding IMPs in the membrane lipid bilayer occurs via hydrophobic protein domains (e.g., hydrophobic helical segments) or covalently-bound lipids or combinations of both and it is energetically costly. Lipids interfacing with IMPs (hereinafter, referred to as annular lipids) provide solvation in the membrane bilayer and exchange with bulk lipids at different rates (Marsh, 2008). Molecular dynamics simulations (MDS) allow an increasingly faithful representation of the lipid bilayer environment sheathing IMPs at the molecular scale (Marrink et al., 2019). These studies have corroborated theoretical predictions and provided support for experimental evidence, together suggesting considerable perturbation of the lipid bilayer thickness, lateral packing and mobility in the immediate proximity of IMPs and at significant distance from them (Ebersberger et al., 2020; Marrink et al., 2019; Mouritsen and Bloom, 1984; Niemela et al., 2010; Phillips et al., 2009). MDS studies employing > 60 membrane lipids species modelled in an asymmetric bilayer suggest that different IMPs show recognisable lipid patterns of annular lipids or lipid fingerprints (Corradi et al., 2018). Lipid fingerprints are made of different lipid species and distribution, resulting in altered bilayer thickness and curvature in IMP’s proximity. This is consistent with crystal and cryo-EM structures of IMPs in detergent micelles or embedded in lipid nanodisks that reveal specific lipids in direct contact with the hydrophobic portions (Lee, 2011; Sun et al., 2018). Lipid fingerprints of IMPs should not be too surprising, considering that IMPs’ transmembrane domains or lipidated membrane anchors and their adjacent hydrophilic sequences interacting with lipid polar headgroups, feature considerable structural diversity and high degree of orthologous conservation (Sharpe et al., 2010). Thus, in natural membranes IMPs can sample among a repertoire of several hundred natural phospholipids present in different proportions and exhibiting significant heterogeneity of acyl chains length, saturation and head-groups and diverse sterols (Lorent et al., 2020; Shevchenko and Simons, 2010) that optimise solvation and function. The combinatorial nature of lipid annuli of varying lipid composition and structure (Corradi et al., 2018) can be seen as a probabilistic concept (i.e., probability of annular lipids’ distribution) and has implications for individual and collective behaviour of IMPs. The thermodynamic properties of diverse lipid annuli should impact on IMPs lateral behaviour and help explaining their potential for IMP organisation into dynamic nanoscale protein-lipid condensates in natural membranes. This idea is rooted in lipid bilayer’s thermodynamics explaining liquid-ordered/disordered lipid phase transition, partial demixing/non-homogenous lipid mixing; short-lived “condensed complexes”, and critical fluctuations near miscibility transitions (Brown, 2017; Destainville et al., 2018; Honerkamp-Smith et al., 2009; Katira et al., 2016; Lingwood and Simons, 2010; London, 2005; McConnell and Vrljic, 2003; Mouritsen, 2010; Phillips et al., 2009; Veatch et al., 2007). These basic precepts and models can be extended to lipid annuli (Destainville et al., 2018; Gil et al., 1997; Ingolfsson et al., 2022; Katira et al., 2016; Molotkovsky et al., 2019), implying correlation length as a measure of the annuli characteristic lateral size (Gil et al., 1997). Consistently, MDS studies suggest that a halo of annular lipids of 10-20 nm co-diffuses with a membrane channel (Niemela et al., 2010). This condition can be seen as an individual IMP-lipid nanodomain endowed with unique physical and chemical properties, though not fully phase-separated (Katira et al., 2016), that could influence IMP lateral interactions and foster formation of IMP clusters, optionally stabilised by protein-protein interactions (Destainville et al., 2016; Ingolfsson et al., 2022; Meilhac and Destainville, 2011; Saka et al., 2014). Nanoscopy has provided evidence that at steady state some IMPs occasionally experience lateral confinement (Douglass and Vale, 2005; He and Marguet, 2011; Kusumi et al., 2012) or manifest clustering, though most often induced by external cues (Abankwa et al., 2007; Dustin and Groves, 2012; Varma and Mayor, 1998), lending support to the idea of biomembranes organised into dynamic lipid-protein nanodomains buttressed by actin-regulated cortical membrane proteins (Gowrishankar et al., 2012; Kusumi et al., 2011; Lingwood and Simons, 2010). However, the underlying mechanisms for formation of such nanodomains remain to be supported by experimental evidence.

The regulation of Lck, a Src-family protein tyrosine kinase (PTK) required for T-cell activation, may offer an opportunity to investigate in a biologically relevant setting the functional role of IMPs immediate lipid environment. In unperturbed cells, ≥ 50 % of Lck is enzymatically active (Lck_A_) (Nika et al., 2010) (Fig. **1A**), required for allosterically-induced T-cell antigen receptor (TCR) signalling (Lanz et al., 2021), as Lck_A_ phosphorylates the TCR to initiate T-cell activation (Acuto et al., 2008). Lck is a monotopic IMP, attached by a lipid-anchor to the inner leaflet of the PM (Yurchak and Sefton, 1995). Lck activity is controlled by the cytoplasmic-resident C-terminal Src kinase (Csk), by trans-autophosphorylation and by the IMP tyrosine phosphatase (PTP) CD45, with Csk and CD45 constitutively active (Fig. **1A**) (D’Oro and Ashwell, 1999; Hermiston et al., 2003; McNeill et al., 2007). Phosphorylation of Lck at Y505 by Csk maintains Lck conformationally “closed” and catalytically inactive (Y394/pY505-Lck, (Lck_I_) (Boggon and Eck, 2004) (Fig. **1A**). CD45 dephosphorylates Y505, decreasing intramolecular constraints, yielding Y394/Y505-Lck or primed-Lck (Lck_P_) (Boggon and Eck, 2004) (Fig. **1A**). Lck_P_ is competent to autophosphorylate *in trans* the activation loop of the kinase domain at Y394, inducing major allosteric changes that generate Lck_A_ (pY394/Y505-Lck) (Fig. **1A**). Lck_A_ displays optimal spatial repositioning of residues required for catalysis (Yamaguchi and Hendrickson, 1996) and full substrate accessibility. Lck_A_ is detected in intact cells by Abs specific for pY394 and when isolated from unperturbed T cells it shows the highest *in vitro* kinase activity of all Lck conformers (Nika et al., 2010). CD45 dephosphorylates also pY394 (Fig. **1A**), acting therefore as a key regulator of Lck_A_ (D’Oro and Ashwell, 1999; McNeill et al., 2007). Lck_A_ can be also phosphorylated in part at Y505 (Fig. **1A**), forming pY394-Lck/pY505-Lck, a double-phosphorylated isoform of Lck (Lck_ADP_) (Nika et al., 2010) that cannot close (Sun et al., 1998) and is enzymatically active similar to Lck_A_ (Nika et al., 2010). Lck_ADP_ generation, cellular localization and role remain unknown. In live cells, pharmacological inhibition of Lck activity drastically reduces Lck_A_, due to dephosphorylation by CD45 (Nika et al., 2010) (and this work). Remarkably, CD45 is in stoichiometric excess over Lck (Hui and Vale, 2014; Nika et al., 2010), making the mechanism of Lck_A_ formation and upkeep intriguing. Lck experiences partial dynamic confinement, (Douglass and Vale, 2005) and it is partially extracted in detergent-resistant membranes (DRMs) (Janes et al., 2000), suggesting that Lck can be dynamically entrapped within a membrane nanodomain (or lipid raft) (Douglass and Vale, 2005). CD45 might have limited access to such nanodomains, favouring therefore Lck_A_ accumulation.

To investigate the molecular mechanism of Lck_A_ upkeep, we genetically swapped Lck membrane anchor with structurally divergent ones borrowed from other IMPs, including single-pass helical anchors of bitopic IMPs and implying considerable alteration of Lck immediate lipids environment (Marrink et al., 2019). We monitored by flow-cytometry Lck_A_ (i.e., at the single-cell and cell-population level) for quantitatively recording the efficacy of the underlying phosphorylation and dephosphorylation mediated by Lck with itself and with CD45. This allowed to claim the occurrence of the highest spatial resolution for IMPs lateral interactions (i.e., a few nm apart required for a chemical reaction). The data showed high tolerance for Lck_A_ upkeep (i.e., of its lateral interactions) to substantial changes in Lck immediate lipid environment. This did not mean that direct protein-protein interaction was the only driver for steady Lck_A_, as small but significant differences in Lck_A_ were recorded with different anchors and Lck_A_ was drastically reduced when endowed with the membrane anchor of CD45, its negative regulator. We discuss how our data identify a modulatory role of annular lipids in the lateral distribution of Lck and its extension to other IMPs.

**Figure 1.**
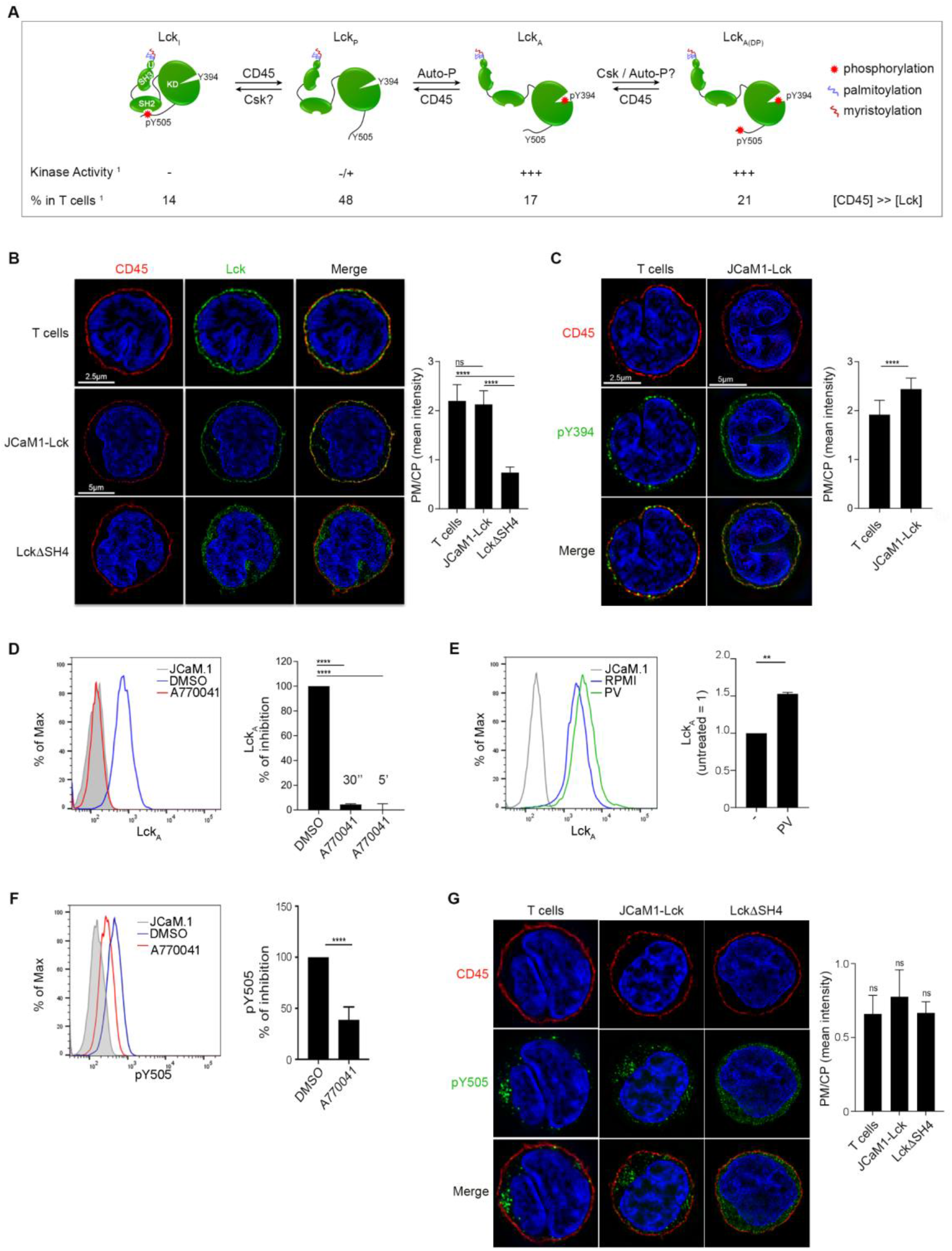
Dynamic maintenance of the Lck_A_ pool. (**A**) Schematic representation of Lck isoforms’ generation, kinase activity and spatial distribution in T cells. From left to right: Inactive Lck (Lck_I_), Primed Lck (Lck_P_), Active Lck (Lck_A_), Active-Double Phosphorylated Lck (Lck_A(DP)_). Symbols represent: tyrosine phosphorylation (red star), palmitoylation (violet curve) and myristoilation (dark-red curve). > indicates that CD45 is in stoichiometric excess over PM-resident Lck. (**B**) **Left**, 3D-SIM of Lck (green) in CD4^+^ T cells or JCaM.1 cells expressing Lck or LckΔSH4. Scale bars: 5 or 2.5 μm. Membrane marker CD45 (red), Nuclear marker DAPI (blue). **Right**, quantitative analysis of Lck distribution at the Plasma Membrane (PM) and Cytoplasm (CP). Bars represent PM/CP ratio of Lck and LckΔSH4. Error bars: SD for n ≥ 10 cells from 3 or more independent experiments, unpaired *t*-test: *p* > 0.5 (non-significant, ns, CD4^+^ T cells vs. JCaM.1-Lck); *p* < 0.0001 (CD4^+^ T cells vs. LckΔSH4). (**C**) **Left**, 3D-SIM imaging of pY394-Lck (green) in CD4^+^ T cells or in JCaM.1 expressing Lck. Scale bars: 2.5 μm and 5 μm, respectively. Membrane marker CD45 (red), Nuclear marker DAPI (blue). **Right**, PM/CP ratio of pY394 in CD4^+^ T cells or in JCaM.1 expressing Lck. Error bars: SD for n ≥ 10 cells from 3 or more independent experiments, unpaired *t*-test *p* < 0.0001. (**D**) **Left**, representative Flow CytoMetry (FCM) of Lck_A_ in Jurkat cells treated (red line) with 2 μM A770041 or carrier (DMSO, blue line) for 30” or 5’ at 37 oC. JCaM.1 (grey) were used as a negative control to set pY416 antibody (Ab) background. **Right**, mean ± SD of Lck_A_ (% of inhibition), n = 3, unpaired *t*-test *p* < 0.0001. (**E**) **Left**, representative FCM of Lck_A_ in Jurkat cells reacted (green line) or not (blue line) with catalase-treated pervanadate (PV) for 1’ at 37 oC. JCaM.1 (grey) were used as a negative control to set pY416 Ab background. **Right**, mean ± SEM of Lck_A_ n = 2, unpaired *t*-test *p* < 0.01. (**F**) **Left**, representative FCM of pY505-Lck in Jurkat cells treated (red line) with 5 μM A770041 or carrier (DMSO, blue line) for 5’ at 37 oC. JCaM.1 (grey) were used as a negative control to set pY505-Lck Ab background. **Right**, mean ± SD of Lck_A_ (% of inhibition), n = 4, unpaired *t*-test *p* < 0.0001. (**G**) **Left**, 3D-SIM imaging of pY505-Lck (green) in CD4^+^ T cells or in JCaM.1 expressing Lck or LckΔSH4. Scale bars: 2.5 μm and 5 μm, respectively. Membrane marker CD45 (red), Nuclear marker DAPI (blue). **Right**, PM/CP ratio of pY505 in CD4^+^ T cells or in JCaM.1 expressing Lck or LckΔSH4. Error bars: SD for n ≥ 10 cells from 3 or more independent experiments, *p* > 0.5 (non-significant, ns).

## Results

### Dynamic maintenance of steady Lck_A_

To validate Lck as a suitable probe for investigating the role of annular lipids in IMPs’ lateral interactions, we assessed the spatiotemporal backdrop for the generation and maintenance of Lck_A,_ schematised in Fig. **1A**. Primary T cells and Jurkat cells, a convenient T-cell surrogate model, were examined for a quantitative assessment of the subcellular distribution of Lck and CD45. We used three-dimensional structured illumination microscopy (3D-SIM) that doubles lateral and axial resolution (i.e., 8-fold in *x, y, z*) and considerably enhances image contrast over conventional fluorescence microscopy (Schermelleh et al., 2008). Staining for CD45 of permeabilised T cells (Fig. **1B**, upper panel) and Lck-deficient Jurkat cells (JCaM.1) expressing Lck (JCaM.1-Lck) (Fig. **1B**, middle panel), showed that CD45 (red) neatly delimits PM-associated regions that with DAPI nuclear staining (blue) emphasises the exiguous cytoplasm (CP) space (see enlargements in Fig. **S1A**). 3D-SIM improved segmentation at regions of interest (ROI) for PM and CP to evaluate preferential distribution for proteins of interest (Fig. **S1A** and Methods). PM/CP ratios > 1 or < 1 reported on the relative preference for PM or CP and allowed to estimate the relative fraction of Lck resident in those compartments. Thus, CD45 staining revealed it to be largely confined to the PM (Fig. **1B** and enlargements in Fig. **S1A**). In contrast, PM/CP for Lck (green) in T cells and JCaM1-Lck scored ≈ 2.2-2.3 (Fig. **1B** and negative control Fig. **S1B**, upper panel), indicating that ≈ 70 % of total Lck (Lck_T_) is PM-resident. Intracellular Lck (Fig. **S1A**, upper panel) was more evident in some optical sections, presumably associated with Golgi and recycling compartments (Bouchet et al., 2017). 3D-SIM of LckΔSH4, a mutant lacking the membrane anchor (Fig. **S1A**, lower panel) and therefore not PM-associated was found mostly in the CP with rare PM detection (Fig. **1B**, bottom panel and enlargement in Fig. **S1A**) and scored 0.6 (Fig. **1B**). Exiguous cytoplasmic space, membrane unevenness and potential interactions with the PM of Lck modular domains (Sheng et al., 2016) may explain the non-null score for LckΔSH4, suggesting a slight imprecision in evaluating the proportion of PM-resident Lck. CD45 neat PM staining helped tracing a reliable mask for ImageStream, which has lower resolution than 3D-SIM, but higher statistical robustness (10,000 events recorded). ImageStream found ≈ 80 % of Lck expressed in JCaM.1 to be PM-resident (Fig. **S1C**, see Methods for details), in good agreement with 3D-SIM (Fig. **1B**) and with previous estimates of Lck subcellular distribution (Bouchet et al., 2017). The virtually exclusive PM localisation of CD45 indicates that this compartment is where most of pY505 of Lck_I_ is dephosphorylated to produce Lck_P_, the open conformer able to accomplish autophosphorylation *in trans* of Y394 to generate Lck_A_ (Fig. **1A**) and where CD45 dephosphorylates Lck_A_ at pY394 (D’Oro and Ashwell, 1999; McNeill et al., 2007) to reverse it to Lck_P_ (Fig. **1A**). The net product of this natural condition is a steady pool of PM-resident Lck_A_ in unperturbed T cells, with CD45 continuously igniting, rescinding and refuelling Lck_A_ formation (Fig. **1A**). This dynamic equilibrium is achieved in spite of CD45 being ≈ 10-fold in stoichiometric excess over Lck (Hui and Vale, 2014; Nika et al., 2010), suggesting a non-trivial mechanism occurring at the PM to achieve steady Lck_A_ amounts instead of its annihilation by CD45 (Nika et al., 2010) and see below). To investigate the molecular basis of this dynamic setting, we monitored Lck_A_ in intact cells using an anti-pY416-Src Ab that recognizes pY394. The fidelity in detecting Lck_A_ by anti-pY416 was confirmed by the lack of staining above background of anti-pY416 signal in flow cytometry (FCM) (Fig. **S1D**) and 3D-SIM (Fig. **S1B**) in JCaM.1. Similar results were obtained when a synthetic peptide made of the pY394-containing activation loop sequence of Lck was used to compete for anti-pY416 binding (Fig. **S1E**). Moreover, A770041, a very potent and specific inhibitor of Lck (Stachlewitz et al., 2005) whose IC_50_ of 1.5 nM (see Table **S1**), is ≈ 300-fold and ≈ 250-fold less potent for Fyn (Stachlewitz et al., 2005) and Csk, respectively (Table **S1**), reduced anti-pY416 staining to background level (Fig. **S1E**). Anti-pY416 in 3D-SIM scored a PM/CP ratio of Lck_A_ in T cells and JCaM1-Lck of 2.0 and 2.5 (Fig. **1C**), respectively, indicating that ≈ 66 -71 % of Lck_A_ is PM-resident. Close inspection of several mid-sections found scarce detection of Lck_A_ in the CP (Figs. **1C** and **S1B**), presumably in a recycling compartment (Bouchet et al., 2017), as Lck_A_ is essentially originated at the PM. Instead, LckΔSH4, which is deprived of stable interaction with the PM (and is therefore mostly CP-resident), formed only tiny amounts of Lck_A,_ as detected by FCM (cf. Figs. **S1F** and **S1D**, right panels) but showed abundant pY505 staining (Fig. **S1G**). This is agreement with LckΔSH4 not being targeted at the PM by CD45 and therefore only present as Lck_i_ in the CP, controlled by Csk (Fig. **S1H**). In Cln20 ≈ 50 % of Lck_T_ is Lck_A_ (Nika et al., 2010), in close agreement with recent estimates by a FRET probe (Wan et al., 2019). Since 70-80 % and ≈ 71 % of Lck_T_ and Lck_A_, respectively, are PM-resident, at least 50-60 % of PM-resident Lck should be Lck_A_, which agrees with earlier data (Nika et al., 2010). Lck_A_ regulation was further gauged by monitoring Lck_A_ upon pharmacological inhibition of Lck and CD45 activities with A770041 and catalase-treated pervanadate (PV), respectively. A770041 (2 μM) erased 90 % of pY394 (i.e., Lck_A_) in 30 sec and 100 % after 5 min (Fig. **1D**). Because Cln20 expresses on average 1.2 × 10^5^ Lck_A_ molecules /cell (Nika et al., 2010), this means conversion of ≈ 4 Lck_A_ molecules to Lck_P_ per ms, revealing the power of CD45 to keep the Lck_A_ pool in check (Fig. **1A**) by a fast turnover of Y394 phosphorylation. Consistent with Lck_A_ relentlessly antagonised by CD45, PV rapidly increased Lck_A_ by 50 % (Fig. **1E**) to a ceiling, revealing a PM pool of Lck_P_ being ≈ 50 % of Lck_A_ and ≈ 30 % of total PM resident Lck, in close agreement with previous estimates (Nika et al., 2010). This also suggested that Lck_I_ should be a minor pool of PM-Lck. Surprisingly, when monitoring pY505-Lck upon A770041 treatment, we observed inhibition of ≈ 60 % of pY505-Lck (Fig. **1F**). If Csk was effective in antagonising CD45 at the PM for the phosphorylation of Lck-Y505, A770041, which is exquisitely specific for Lck (Table **S1**), should increase and not decrease Lck-pY505. This result indicated that a considerable proportion of PM-resident pY505-Lck is not generated by Csk but by Lck itself, presumably by trans-autophosphorylation of Lck_A_ at pY505 and forming the double phosphorylated Lck isoform (Lck_ADP_) (Fig. **1A**). This is consistent with *in vitro* or *in cellulo* data that Lck (Hui and Vale, 2014) and Src (Nelson et al., 2020; Sun et al., 1998) can phosphorylate the C-terminal regulatory tyrosine. For evident steric constraints, double phosphorylated Src cannot close (Sun et al., 1998), consistent with Lck_ADP_ featuring *in vitro* kinase activity similar to Lck_A_ (Hui and Vale, 2014; Nika et al., 2010). Lck_ADP_ was therefore considered as part of the Lck_A_ pool as detected by anti-pY416 Ab and its functional role was not explored as beyond the scope of this investigation. Further 3D-SIM analysis revealed that, contrary to Lck_A_, PM/CP ratios of pY505-Lck in T cells and JCaM1-Lck scored 0.7 - 0.8 (Fig. **1G** and see Fig. **S1I** for Ab specificity), respectively, indicating it to be more abundant in CP than PM. Since a sensible proportion of PM-resident pY505-Lck is generated by Lck_A_, the role of Csk in controlling pY505-Lck in this compartment should be resized. Thus, Csk might primarily control Y505-Lck in the CP, keeping in check Lck as Lck_I_ presumably in exocytic compartments *en route* to the PM (Figs. **1G** and **S1H**). Consistently, pY505 PM/CP ratio for Lck was only slightly higher than for LckΔSH4 (Fig. **1G**) and claims that Csk is constitutively recruited to the PM via CBP/PAG to negatively regulate Src PTKs have not been supported by gene knock-out (Dobenecker et al., 2005; Xu et al., 2005). Comprehensively, our imaging and FCM data support a model in which Lck_A_ is generated and maintained at the PM by a ceaseless “Lck cycle”, based on the antagonistic action of CD45 and Lck at Y394 (Fig. **S1H**), whose dynamicity is illustrated by the remarkable dephosphorylation rate of Lck_A_ upon pharmacological inhibition by A770041. Steady maintenance of Lck_A_ may be explained by dynamic segregation of Lck_P_ (favouring auto-phosphorylation by local concentration) and/or Lck_A_ (averting pY394 dephosphorylation by CD45) into liquid ordered (L_o_), phase-separated membrane nanodomains (or rafts). Egress of Lck_A_ from such shelters would expose it to CD45, rapidly reversing it to Lck_P_ to feed Lck_A_ formation. Such Lck raft-philic model may be suggested by Lck being partially found in DRMs (Janes et al., 2000) and lateral diffusion exhibiting short confinement (Douglass and Vale, 2005). In contrast, CD45 undergoes random diffusion over larger distances, partially restricted by transient interaction of CD45 intracellular domain with membrane cortex proteins (Cairo et al., 2010; Douglass and Vale, 2005; Freeman et al., 2016). Alternatively, steady maintenance of Lck_A_ might be explained only by differential kinetics of Lck protein-protein lateral interactions (and catalytic rates) with itself and with CD45, without invoking pre-existing membrane lipid phases.

### Lck_A_ dependence on Lck_T_

To distinguish between these models and seeking for a possible role of annular lipids in Lck_A_ upkeep, we set up an accurate FCM-based assay for quantitatively estimating Lck_A_ output as a function of Lck_T_. We opted for a two-colour FCM-based assay to concomitantly detect Lck_A_ and Lck_T_ on a per cell basis. An anti-Lck Ab (73A5) raised against Lck C-terminal tail, was found to be most adequate for this purpose. 73A5 showed an excellent FCM signal-to-noise ratio and epitope mapping by non-phosphorylated overlapping peptides revealed it to recognise Lck C-terminal end including Y505 (Fig. **S2A**). Treatment by A770041 or PV, both of which can change Y505 phosphorylation and conformers level, left 73A5 reactivity largely unaffected (Fig. **S2B** and **S2C**), indicating that 73A5 does not discriminate among Lck isoforms. 73A5 and anti-pY416 Abs were used at saturating concentrations with negligible effect on signal-noise (see Methods) and no hindrance to one another for Lck binding was observed (Fig. **S2D**). These features should allow to unambiguously quantitate Lck_A_ as a function of Lck_T_ per cell basis and to evaluate this relationship over a considerable Lck_T_ dynamic range. Double staining of Cln20 and FCM provided 2D plots (Fig. **2A**), reporting the dependence of Lck_A_ on Lck_T_ by exploiting Lck normal distribution in the cell population. Lck expression was largely independent of cell-size (Fig. **S2E**), effectively reporting increasing Lck concentration per cell basis. A dense binning was drawn on the 2D plot (Fig. **S2F**, left and middle panel) and the values for Lck_T_ and Lck_A_ extracted for each bin, averaged ± SD and plotted to obtain the best fit by regression analysis (Fig. **S2F**, right panel and see Methods). Lck_A_ formation showed two components: at low Lck_T_ concentration, Lck_A_ fitted a second-order function, whereas at higher Lck_T_ concentration Lck_A_ became linear (Fig. **S2F**). This suggested that the probability of an “active event” (i.e., formation of Lck_A_) depends in general on Lck_T_ but, above a Lck_T_ threshold, it becomes also dependent on Lck_A_. The molecular basis this observation can be inferred from the crystal structure of the unphosphorylated IRAK4 dimer (Ferrao et al., 2014), in which the dimer conformation is highly suggestive of a genuine trans-autophosphorylation reaction, with the phosphorylation site in the activation loop of one IRAK4 monomer correctly positioned for phospho-transfer by its partner. If a pair of Lck_P_ molecules (Lck_P_:Lck_P_) assumes a similar configuration, it is unlikely that two Lck_A_ molecules could form simultaneously, but should rather be produced by two independent reactions (2) and (3) (Fig. **2B**), occurring in series. Moreover, for steric considerations, converting Lck_P_ into Lck_A_ from Lck_P_:Lck_P_ pair can be expected to be less efficient than from a Lck_A_:Lck_P_ pair. If so, then increase of Lck_A_:Lck_P_ pairs (i.e., in cells expressing more Lck_T_, Fig. **2B**) should increase the yield of Lck_A_. To test the validity of these assumptions, numerical simulations were carried out by allowing the probability of converting Lck_P_ into Lck_A_ from reaction (2) P_PA_ and (3) P_AA_ (Fig. **2B**) to vary between 0.1 and 1.00 (with incremental steps of 0.05) (Fig. **2C**, and see Methods for details of the modelling). The best fit of the numerical simulation to the experimental data was obtained for P_PA_ and P_AA_ of 0.1 and 0.3, respectively (Fig. **2C**, insert). This result agrees with Lck_A_ being produced more efficiently by a Lck_A_:Lck_P_ pair (P_AA_ > P_PA_), supporting a two-step reaction scheme in Fig. **2B** and helps explaining a mechanism of Src kinases trans-autophosphorylation. Fig. **2D** resumes qualitatively the dynamic equilibrium suggested by our data. Independently of the exact structural details of transactivation for Lck pairs and the actual kinetic parameters of the underlying reactions, our data provided a sound quantitative basis for asking whether perturbing Lck hydrophobic anchor (i.e., the immediate lipid environment) modifies Lck_A_ output.

**Figure 2.**
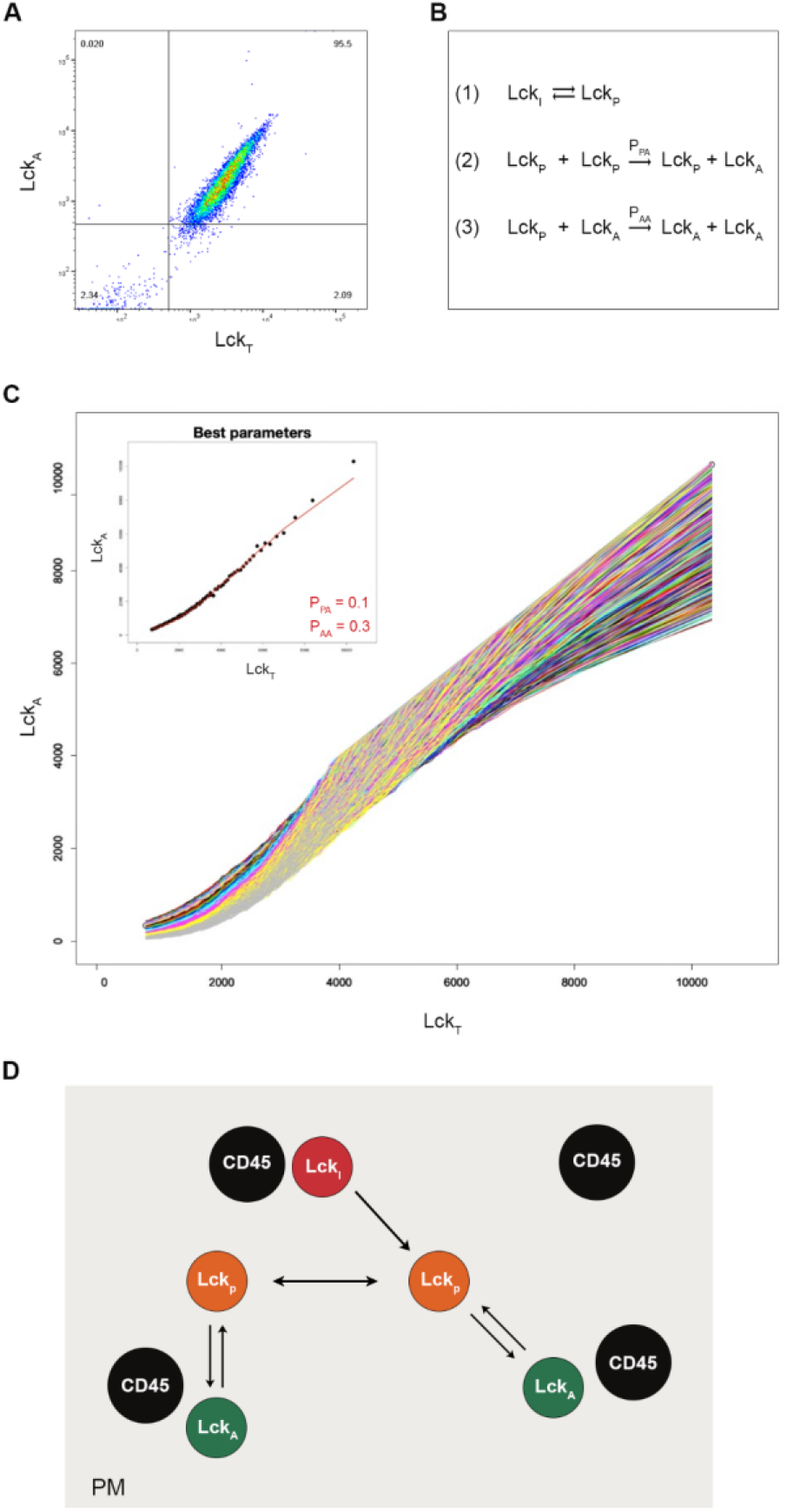
Lck_A_ dependence on Lck_T_. (**A**) Representative FCM 2D plot of Cln20 concomitantly stained for Lck_A_ and Lck_T_. (**B**) Reactions considered to generate the probabilistic model for Lck_A_ formation as a function of Lck_T_. Lck_I_, Lck_P_ and Lck_A_ indicate the inactive (closed), primed and active Lck isoforms, respectively. P_PA_ and P_AA_ are the probabilities of generating Lck_A_ from the reactions (Lck_P_ + Lck_P_) and (Lck_P_ + Lck_A_), respectively. See Main Text and Methods for further details on the rational basis of the model. (**C**) The increase of Lck_A_ as a function of Lck_T_ as obtained by changing the two probabilities of the reactions (2) and (3) showed in **B**. The best agreement with experimental data with the associated probability values are reported in the inset. P_PA_ and P_AA_ are the probabilities of generating Lck_A_ from the reactions (Lck_P_ + Lck_P_) and (Lck_P_ + Lck_A_), respectively. (**D**) Schematic representation of the “Lck cycle” at the PM where Lck_A_ is generated and maintained by the antagonism between CD45 and Lck for Y394 phosphorylation. At the PM, Lck_I_ is rapidly dephosphorylated at Y505 by CD45 and converted in Lck_P_. Lck_P_ in turn generates Lck_A_ by two independent reactions: Lck_P_:Lck_P_ or Lck_A_:Lck_P_ pair, as shown in **B**. Lck_A_ can either be dephosphorylated by CD45 or accumulates at the PM.

### Subcellular distribution of Lck with non-native membrane anchors

Together, myristoylation and di-palmitoylation of LckSH4 (Fig. **3A**) provide firm attachment to the inner leaflet of the PM (Yurchak and Sefton, 1995). Palmitoylation is thought to partition Lck within phase-separated lipid nanodmains (Lorent et al., 2017), a property that should contribute to concentration and sheltering from CD45 and to ensure Lck_A_ steady maintenance. LckSH4 is eleven amino acid-long and devoid of secondary structure, away from folded Lck SH domains. As such, LckSH4 is unlikely to have a critical influence on Lck allosteric regulation and catalytic activity. However, being the predominant Lck lipid bilayer-interacting moiety, LckSH4 should play a major role in defining Lck immediate lipid environment. We surmised that swapping LckSH4 with structurally diverse IMPs’ membrane anchors, including removing palmitoylation (Lorent et al., 2017), should inform about the role of Lck immediate lipid in Lck_A_ formation and maintenance. Therefore, Lck lacking SH4 (LckΔSH4) was fused to membrane anchors of considerably disparate physicochemical properties (Fig. **3A** and Table **S2**). They included the SH4 domain of Src (SrcSH4), which is myristoylated but not palmitoylated and, contrary to LckSH4, contains several basic residues (Table **S2**), which may mediate interactions with negatively-charged phospholipids (Marrink et al., 2019). Moreover, we chose helical TM anchors of the scaffold protein LAT and of the co-receptor CD4, both featuring two palmitoylation sites and a palmitoylation-defective CD4 TM helix mutant (CD4C/S). Palmitoylation-defective LAT is not expressed at the PM (Tanimura et al., 2006) and the effect of TMD LAT palmitoylation could not be tested. These helical anchors diverged for amino acid composition, sequence, length and membrane-juxtaposed segments (Table **S2**). Consequently, they should confer to Lck annular lipids of diverse composition and topological arrangement as compared to native Lck (Marrink et al., 2019; Corradi et al., 2018; Lee, 2011). Moreover, none of those TMDs has been reported to contribute to dimer formation (Parrish et al., 2015; Sherman et al., 2011), making it unlikely that they favour Lck pairing. The three residues-long extra-cellular sequence of LAT was added to each helical anchor to facilitate similar expression of the Lck chimeras. To minimise potential phenotypic drift of cell cultures, we conditionally expressed Lck or chimeras in JCaM.1 by dox induction for 16 h. All chimeras were expressed similarly to Lck (Fig. **3B**), with only SrcSH4-Lck expressing about twice as much and all cell lines maintained identical amounts of endogenous CD45 (Fig. **S3A**). Inspection of PM/CP ratios determined by 3D-SIM for LAT-Lck, CD4-Lck and CD4C/S-Lck chimeras (Fig. **3C**) indicated them to be very similar to native Lck. Only SrcSH4-Lck showed a PM/CP of about 1.00 (i.e., even PM and CP distribution), perhaps reflecting Src higher propensity to localise in recycling membranes (Sandilands and Frame, 2008). However, SrcSH4-Lck reduction at the PM should be compensated by its total higher expression (Fig. **3B**) and results in PM-resident SrcSH4-Lck absolute amount similar to the other chimeras. Thus, all non-native membrane anchors conferred PM residency similar to native Lck, guaranteeing a fair comparison of their capacity to form Lck_A_.

**Figure 3.**
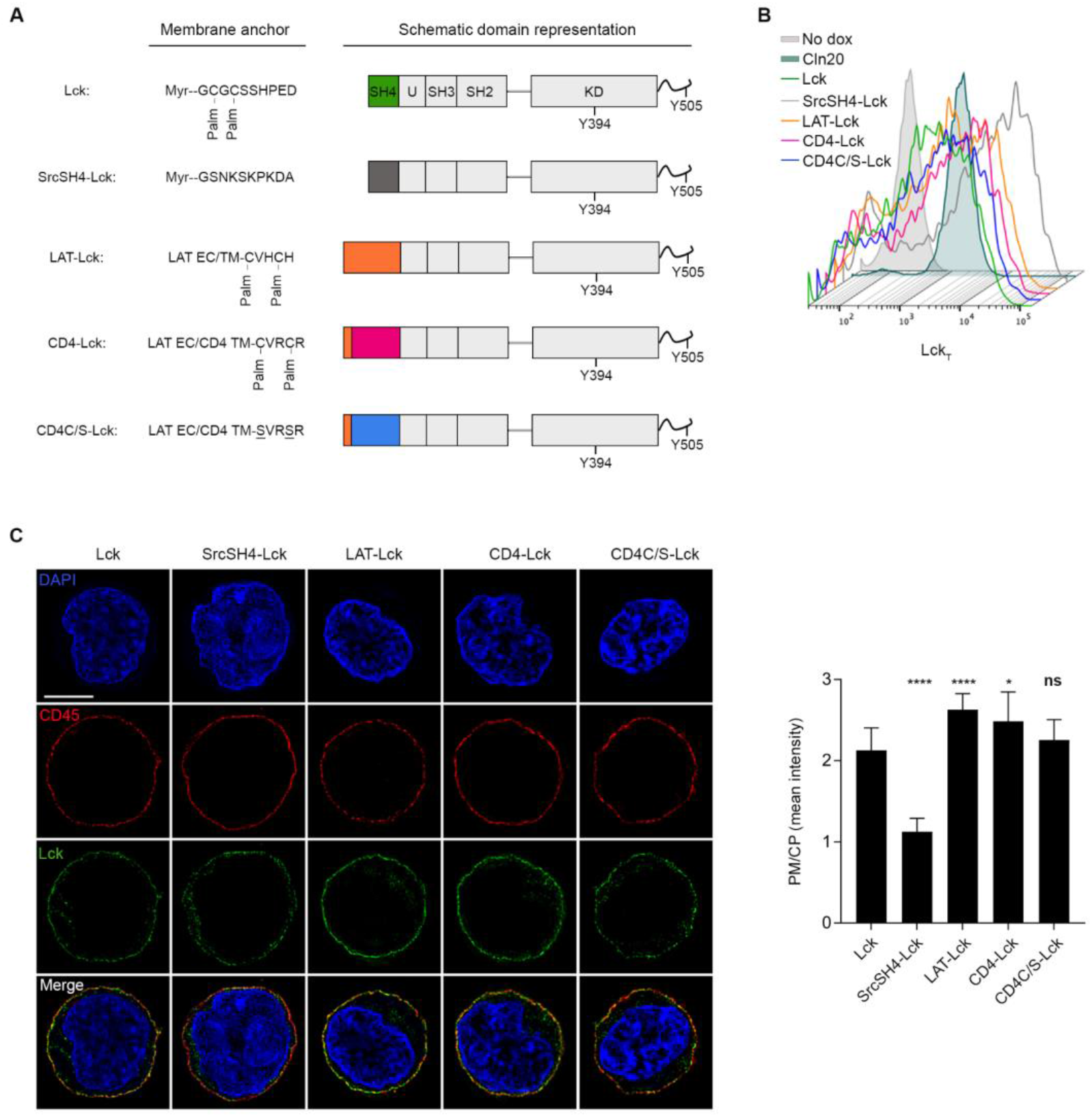
Subcellular distribution of Lck with non-native membrane anchors. (**A**) Schematic representation of Lck or Lck-chimeras employed in this investigation. (**B**) Representative FCM of Lck_T_ in Cln20 and JCaM.1 cells conditionally expressing Lck or the indicated Lck-chimeras by 16-18 h of doxycycline (dox) induction. Non-dox-induced cells have been used to assess Ab background. (**C**) **Left**, representative 3D-SIM imaging of Lck (green) in JCaM.1 cells expressing the constructs showed in **A**. Scale bars: 5 μm. Membrane marker CD45 (red), Nuclear marker DAPI (blue). **Right**, PM/CP for Lck of the indicated Lck constructs. Error bars: SD for n ≥ 10 cells from 3 or more independent experiments, unpaired *t*-test: *p* < 0.0001 (Lck vs. SrcSH4-Lck); *p* < 0.0001 (Lck vs. LAT-Lck); *p* < 0.05 (Lck vs. CD4-Lck); *p* > 0.05; (non-significant, ns, Lck vs. CD4C/S-Lck).

### Moderate impact of different membrane anchors on Lck_A_ formation

Lck and chimeras were conditionally expressed by dox and stained for Lck_T_ and Lck_A_ for FCM analysis to obtain 2D plots (Fig. **4A**). A cell tracer was used to augment data robustness by mixing before dox-induction two chimera-expressing cells together with native Lck-expressing cells (used as standard), followed by FCM (Fig. **4A** and see Methods for details). A gate delimiting Lck_T_ expression in Cln20 was applied to the 2D plots to restrict the analysis of Lck_A_ formation to a physiological range of Lck expression (blue box superimposed to each FCM 2D plot in Fig. **4A** and **S4A**). This was justified by Cln20 expressing about 4-5 times higher Lck than T cells (Nika et al., 2010), but having twice the diameter of a T cell (Fig. **1B**) (i.e., 4 times larger cell surface) and Cln20 and T cells showing similar PM/CP for Lck (Fig. **1B**). The distribution of dox-induced Lck_T_ (and Lck_A_) was not influenced by cell size (Fig. **S4B**), excluding constant Lck_T_ concentration per cell. Steady Lck_A_ for every chimera continued to increase with a similar dependency on Lck_T_, even at Lck_T_ expression ≥ 5-10-fold higher than in Cln20. This indicated a considerable reservoir of PM properties and CD45 enzymatic capacity to allow Lck_A_ formation and control. This relatively ample dynamic range of Lck_A_ generation was unlikely to depend on a dedicated membrane scaffold protein, which may be expected to be a limiting factor. FCM 2D plots were binned and data within the Cln20 range extracted as in Fig. **S2F** (Fig. **4B**, left panels) and subject to regression analysis for best fitting (Fig. **4B**, right panels). Surprisingly, the data showed only small differences in Lck_A_ formation by SrcSH4-Lck, LAT-Lck (Fig. **4B** upper panels), CD4-Lck and CD4C/S-Lck (Fig. **4B** bottom panel), as compared to native Lck. Regression analysis indicated that all the curves of the chimeras were not overlapping (Fig. **4B**, right panels), suggesting that such small differences in Lck_A_ were significant. Similar results were obtained by plotting Lck_A_ normalised to Lck_T_ for each bean (Lck_A_/Lck_T_ vs. Lck_T_ plots in Fig. **S4C**). These plots better capture the two regimens of Lck_A_ yield at low and high Lck_T_, as observed for Cln20. Predictably, LckΔSH4 showed severely reduced Lck_A_ (Figs. **S1E, 4C** and **S4D**), despite being expressed at higher amounts than Lck (Fig. **S4E**) and for equal CD45 expression (Fig. **S4F**), consistent with LckΔSH4 being not PM-anchored and therefore escaping CD45 regulation (Fig. **S1H**) and representing therefore the lowest capacity of Lck to generate Lck_A_ in our experimental system. The behaviour of the Lck chimeras was unexpected in view of the substantial physicochemical divergence of the hydrophobic anchors. In particular, palmitoylation was found not to be essential, nor providing an advantage for Lck_A_ generation. Rather, LAT-Lck and CD4-Lck performed slightly worse than Src-Lck and CD4C/S-Lck that are not palmitoylated (Fig. **4B**), both of which showed a small but significant improvement of Lck_A_ steady formation compared to native Lck (Fig. **4B**). Because four of four chemically diverse membrane anchors did not severely compromise steady Lck_A_ level, one could conclude that direct protein-protein interaction (Lck:Lck and Lck:CD45) are dominant and Lck and CD45 immediate lipid environment may have a modulatory effect that tunes steady Lck_A_. Alternatively, all membrane anchors could confer to Lck very similar lateral behaviour with similar stalling within a different phase-separated nanodomain. Being less likely (see Discussion), the latter hypothesis warranted to test an alternative explanation for our observations.

**Figure 4.**
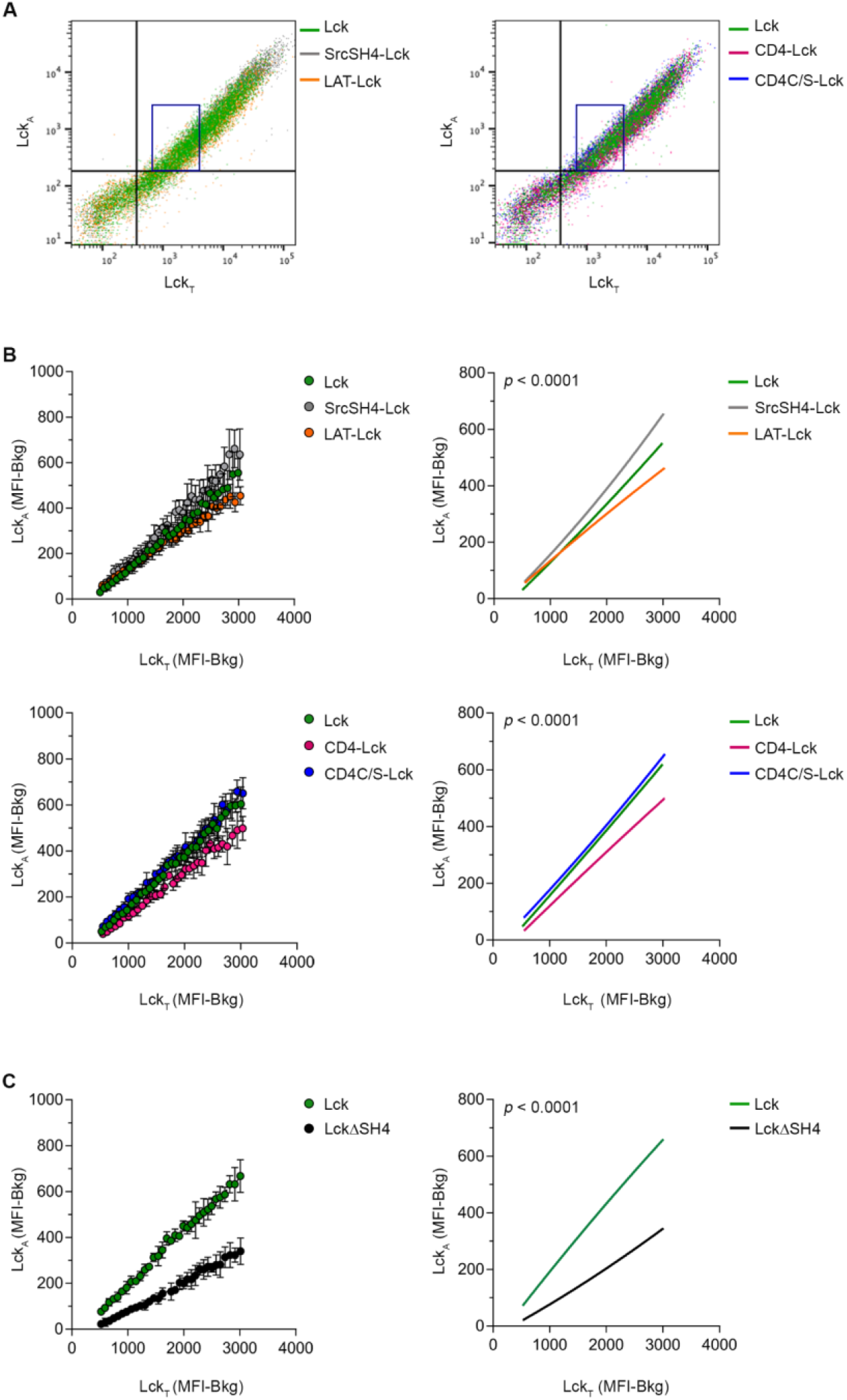
Moderate impact of different membrane anchors on Lck_A_ formation. (**A**) Representative FCM 2D plot of JCaM.1 expressing Lck or Lck-chimeras concomitantly stained for Lck_A_ and Lck_T_. The blue box represents the limits for Lck_A_ and Lck_T_ of Cln20. **Left**, FCM 2D plot of JCaM.1 expressing Lck (green), SrcSH4-Lck (grey) or LAT-Lck (orange). **Right**, FCM 2D plot of JCaM.1 expressing Lck (green), CD4-Lck (magenta), CD4C/S-Lck (blue). (**B**) Lck_A_ formation depending on Lck_T_ of JCaM.1 expressing Lck (green), SrcSH4-Lck (grey), LAT-Lck (orange), CD4-Lck (magenta), CD4C/S-Lck (blue). The indicated cells were labelled or not with two different concentrations of CellTrace violet, mixed 1:1:1, induced for Lck expression by dox and, 16-18 h after, concomitantly analysed by FACS for Lck_A_ and Lck_T_. A dense binning within a physiological concentration range of Lck_T_ set on Cln20 was applied and the values of the geometric median for Lck_A_ and Lck_T_ in each bin, were extracted. **Upper left**, 2D plot of the extracted experimental values of the geometric median for Lck_A_ and Lck_T_ in each bin in JCaM.1 cells expressing Lck or the indicated Lck chimera. **Upper right**, non-linear regression fit of Lck_A_ (MFI-Bkg) vs. Lck_T_ (MFI-Bkg), n = 3, R^2^ = 0.99 (Lck), 0.99 (SrcSH4-Lck), 0.99 (LAT-Lck); F-test *p* < 0.0001. **Bottom left**, 2D plot of the extracted experimental values of the geometric median for Lck_A_ and Lck_T_ in each bin in JCaM.1 cells expressing Lck or the indicated Lck chimera. **Bottom right**, non-linear regression fit of Lck_A_ (MFI-Bkg) vs. Lck_T_ (MFI-Bkg), n = 3, R ^2^= 0.99 (Lck), 0.99 (CD4-Lck), 0.99 (CD4C/S-Lck); F-test *p* < 0.0001. See also Fig. **S4C**. (**C**) Lck_A_ formation depending on Lck_T_ of JCaM.1 expressing Lck (green) or LckΔSH4 (black). Cells were treated and data processed as in **B. Left**, 2D plot of the extracted experimental values of the geometric median for Lck_A_ and Lck_T_ in each bin in JCaM.1 cells expressing Lck or or LckΔSH4. **Right**, non-linear regression fit of Lck_A_ (MFI-Bkg) vs. Lck_T_ (MFI-Bkg), n = 3, R^2^ = 0.99 (Lck), 0.99 (LckΔSH4); F-test *p* < 0.0001. See also Fig. **S4D**.

### How Lck membrane anchor impacts on lateral interactions

To provide a comprehensive explanation for our puzzling data, we considered the possibility that identity and diversity in membrane anchors (i.e., annular lipids) favours IMP lateral proximity or distancing, respectively. It has been theoretically predicted (Mouritsen and Bloom, 1984; Marsh, 2008; Phillips et al., 2009) and suggested by recent MDS studies (Corradi et al., 2018; Marrink et al., 2019) that solvation of IMPs in the lipid bilayer alters the arrangement of the (bulk) lipid bilayer in the IMP proximity. This condition entails quantitative and qualitative differences in composition of annular lipids with respect to the average bulk lipids composition. Differences in annular lipid arrangement and lipid miscibility can generate energy barriers between IMPs comparable to or larger than the thermal energy (Destainville and Foret, 2008; Gil et al., 1997; Reynwar and Deserno, 2008) that can be theoretically estimated to be of few Kcal/mole and affect the likelihood of IMP lateral proximity, without however forbidding it. In contrast, these energy barriers should be much lower or even vanishingly small for identical lipid environments (i.e., identical hydrophobic bilayer anchors). Model simulations agree with this mechanism (Destainville and Foret, 2008; Meilhac and Destainville, 2011; Reynwar and Deserno, 2008), which does not necessarily require IMPs’ trapping in a phase-separated lipid nanodomains. Accordingly, independently of direct protein-protein interaction, the frequency of Lck lateral proximity with itself should be favoured, but it should be less so between Lck and CD45 that have highly different hydrophobic anchors (hence, annular lipids). If so, each Lck chimera can generate Lck_A_ similarly to native Lck, but manifest lateral proximity to CD45 dissimilar from native Lck, resulting in small differences (even of different sign) in the efficacy to upkeep Lck_A_ (Fig. **4**). One prediction of this conjecture is that Lck endowed with the CD45 TMD (CD45-Lck) (Fig. **5A**) should exhibit trans-autophosphorylation capacity (i.e., Lck_A_ generation) similar to native Lck. However, CD45-Lck should have a higher likelihood of dynamic proximity to endogenous CD45 and consequently experience reduction or annihilation of steady Lck_A_. To test this prediction, LckΔSH4 was fused to CD45 helical TMD (CD45-Lck) (Fig. **5A** and Table **S2**) and conditionally expressed in JCaM.1 at similar levels as native Lck (Fig. **S5A**). 3D-SIM for CD45-Lck showed a TM/CP ratio of 1.7 (Fig. **5B**), only slightly lower than Lck (i.e., 63 % vs. 68 % PM-resident for CD45-Lck and Lck, respectively). In agreement with the above prediction, CD45-Lck yielded drastically lower Lck_A_ formation than native Lck (and other Lck chimeras) and was virtually indistinguishable from LckΔSH4 (Fig. **5C** and **S5B**). Expression of endogenous CD45 was identical to cells expressing native Lck (Fig. **S5C**), excluding that changes in CD45 explained Lck_A_ reduction. To test whether the striking reduction of Lck_A_ was due to accrued capacity of endogenous CD45 to dephosphorylate CD45-Lck_A_, we acutely inhibited CD45 enzymatic activity by moderate concentration of PV. The data showed that PV induced rapid recovery of CD45-Lck_A_ (Fig. **5D**), and are schematised in Fig. **5E**. Thus, CD45-Lck has no impediment of accomplishing trans-autophosphorylation but experiences a dephosphorylation rate of pY394 by endogenous CD45 considerably higher than native Lck. Lck_A_ increment by PV for Lck and CD45-Lck above their respective basal Lck_A_ values was very similar (Fig. **5D**), further excluding alterations of CD45-Lck trans-autophosphorylation ability and indicating that CD45-Lck forms Lck_A_ with similar capacity as native Lck. Note that PV treatment of cells expressing LckΔSH4 showed poor recovery of Lck_A_ (Fig. **5D**), indicating different causes for reduced Lck_A_ of CD45-Lck and LckΔSH4, namely, poor trans-autophosphorylation capacity of the latter because it lacks membrane docking. Thus, an apparently simple rule of lateral proximity and distancing driven by annular lipids can explain our data (see Fig. **5E**), a mechanism not necessitating membrane nanodomains, at least in their original conception.

**Figure 5.**
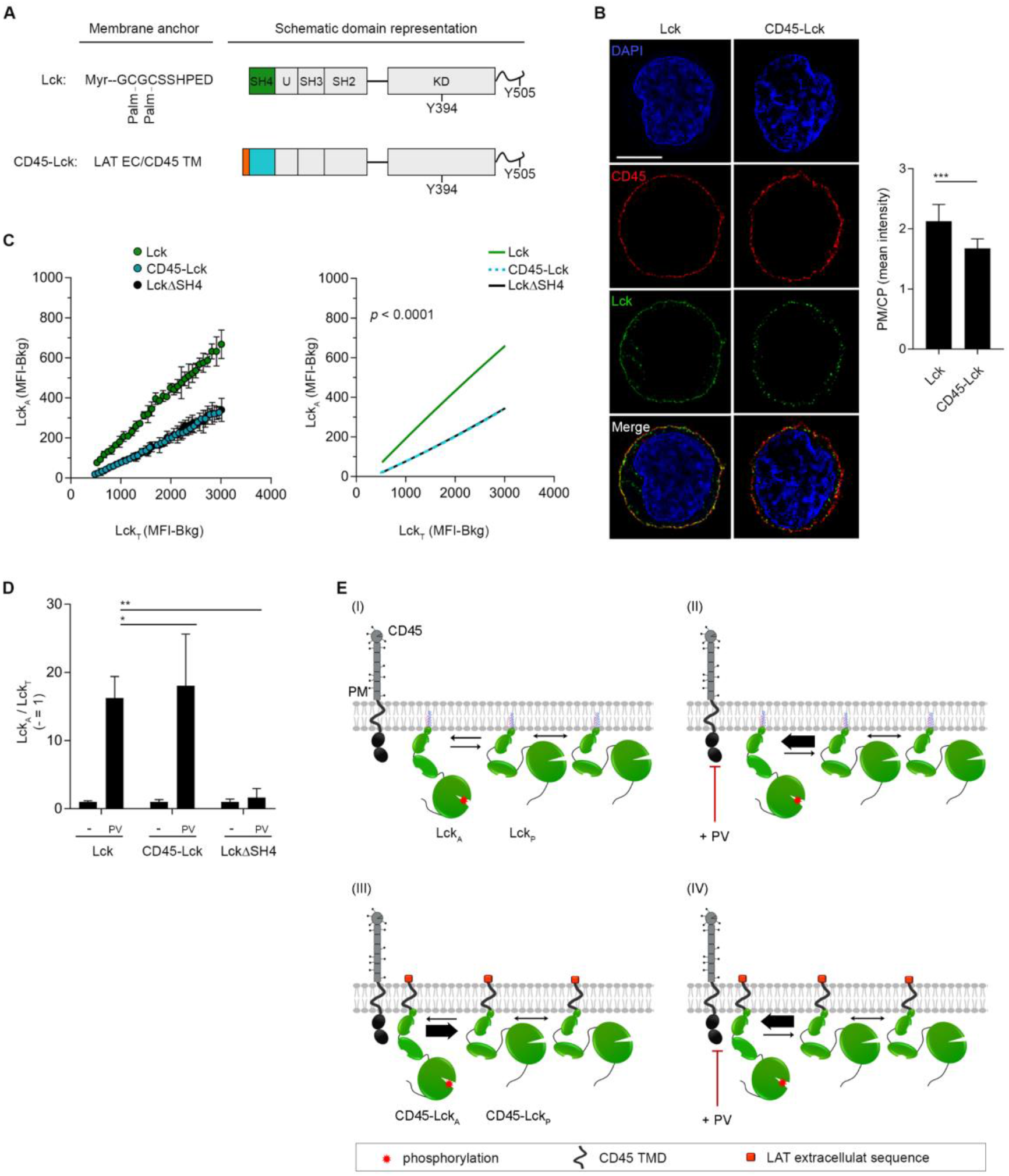
How IMPs lipid anchor can impact on lateral interaction. (**A**) Schematic representation of CD45-Lck chimera compared to Lck. (**B**) **Left**, representative 3D-SIM imaging of Lck (green) in JCaM.1 cells expressing Lck or CD45-Lck. Scale bars: 5 μm. Membrane marker CD45 (red), Nuclear marker DAPI (blue). **Right**, PM/CP for Lck of the indicated Lck constructs. Error bars: SD for n ≥ 10 cells from 3 or more independent experiments. (**C**) Lck_A_ formation depending on Lck_T_ of JCaM.1 expressing Lck (green), CD45-Lck (cyan) or LckΔSH4 (black). The indicated cells were labelled or not with two different concentrations of CellTrace violet, mixed 1:1:1, induced for Lck expression by dox and, 16-18 h after, concomitantly analysed by FACS for Lck_A_ and Lck_T_. A dense binning within a physiological concentration range of Lck_T_ set on Cln20 (blue box) was applied and the values of the geometric median for Lck_A_ and Lck_T_ in each bin, were extracted. **Left**, 2D plot of the extracted experimental values of the geometric median for Lck_A_ and Lck_T_ in each bin in JCaM.1 cells expressing Lck or the indicated Lck chimera or mutant. **Right**, Non-linear regression fit of Lck_A_ (MFI-Bkg) vs. Lck_T_ (MFI-Bkg), n = 3, R^2^ = 0.99 (Lck), 0.99 (CD45-Lck), 0.99 (LckΔSH4); F-test *p* < 0.0001. See also Fig. **S5B**. (**D**) Increase of Lck_A_ of JCaM.1 expressing Lck, CD45-Lck or LckΔSH4 treated or not with 100 μM PV for 3’ at 37 oC. Bars indicate mean ± SEM of Lck_A_/Lck_T_, n = 2, unpaired *t*-test *p* < 0.05 (Lck vs. CD45-Lck) and *p* < 0.01 (Lck vs. LckΔSH4). (**E**) Schematic representation of CD45 dephosphorylation ability of Lck_A_ for native Lck or CD45-Lck. (I) Lck_A_ generated by trans-auto phosphorylation at the PM, is partially reverted to Lck_P_ by CD45. (II) Inhibiting CD45 enzymatic activity by pervanadate (PV) results in higher level of Lck_A_. (III) CD45-Lck chimera shares the same anchoring of the CD45 phosphatase and experiences augmented proximity to CD45 resulting in dramatic reduction of Lck_A_ (thicker arrow of Lck_A_ reversion to Lck_p_). Note that Y394 trans-autophosphorylation should remain intact. (IV) PV rescues Lck_A_ upkeep to wild-type level indicating that CD45-Lck can form Lck_A_ with similar capacity as native Lck.

## Discussion

We found that the steady maintenance of Lck_A_ at the PM of T cells depends on a previously unrecognised role of Lck membrane anchor in modulating its lateral behaviour, a notion that may be generalised to other IMPs. A key finding was that the replacement of Lck native membrane anchor by the TMD of CD45 massively inhibited of Lck_A_, a condition that could be swiftly reversed to Lck_A_ wild-type level upon inhibition of CD45 enzymatic activity (Fig. **5E**). The identity of membrane anchor explains the augmented lateral proximity of Lck to CD45, reminiscent of elegant experiments reported two decades ago. Thomas and co-workers (He et al., 2002) observed that Lck tyrosine phosphorylation (and TCR-proximal signalling) was vigorously inhibited in T cells expressing CD45 intracellular domain anchored to PM via Lck-SH4. Tsien and co-workers (Zacharias et al., 2002) found that mutated GFP and YFP (mGFP and mYEF), which cannot form dimers in solution, exhibited FRET (i.e., requiring proximity of a few nm) when anchored to the PM via the same membrane anchor, being either dual-acylation or prenylation. However, FRET was markedly reduced when mGFP and mYEF were dual-acylated and prenylated, respectively, and vice versa. These studies and our data indicate that membrane anchor sameness or variance can confer to IMPs lateral proximity or remoteness, respectively. Both early studies concluded that each lipidated membrane anchors conferred bestowed confinement (i.e., concentration) in different lipid raft, favouring therefore proximity or remoteness (He et al., 2002; Zacharias et al., 2002). As discussed below, our experimental evidence as well as current notions on annular lipids warrant alternative explanations.

Lck diffuses randomly interrupted by brief confinement with no immobilisation, thought to be the result of occasional interaction with lipid rafts (Douglass and Vale, 2005). GFP coupled to Lck-SH4 has similar confined diffusion (Lommerse et al., 2006), suggesting that Lck anchor alone confers this type of lateral behaviour. CD45 diffusion is passive and random with occasional halts due to a specific interaction of CD45 intracellular moiety with spectrin-ankyrin (Cairo et al., 2010; Douglass and Vale, 2005; Freeman et al., 2016). Thus, CD45 TMD should not confer to CD45-Lck trapping into a lipid raft as Lck-SH4 anchor is thought to do and CD45-Lck should experience random diffusion. This makes unlikely that Lck_A_ annihilation is due to trapping of CD45-Lck and native CD45 in the same lipid raft, as earlier explanation would have suggested (Zacharias et al., 2002). Pharmacological inhibition of CD45 immediately rescued Lck_A_ generation by CD45-Lck similar to native Lck (Fig. **5E**), indicating that a lipidated membrane anchor is not mandatory for Lck_A_ generation nor its trapping in a conventional lipid raft. Indeed, as shown here, effective Lck trans-autophosphorylation at Y394 occurred with a disparate choice of membrane anchors made of myristylation only or hydrophobic helical TMDs, including CD45 TMD. Significantly, Lck_A_ generation did not require palmitoylation, considered as a landmark for IMPs partitioning into L_o_ nanodomain (Lorent et al., 2017) and absence of palmitoylation slightly improved Lck_A_ formation. Quantitative considerations make also unlikely that disparate lipid nanodomain subsets exist for each and every membrane anchor tested and are able to comparably trap Lck and limit access to CD45, a possibility further weakened by the ≈ two-order magnitude scalability of Lck_A_ formation by Lck and Lck chimeras (Fig. **4A**).

From a theoretical perspective, the possibility of enhanced protein-protein encounter in the absence of pre-defined rafts or protein long-lived nanodomains, while remaining related to lipid mixture demixing, was pioneered 25 years ago by Mouritsen and coworkers (Gil et al., 1998; Gil et al., 1997). The mechanism relies on the “wetting” of IMPs by lipid annuli, the lateral extension of which is set by the correlation length of the lipid mixture. This correlation length becomes very large when getting close to a second-order demixing transition and can vary from a few to several lipid shells around each IMP. More precisely, the study of critical phenomena states that when a lipid mixture is close to a second-order transition, extended density fluctuations appear in the medium, whose typical size is set by the correlation length (Destainville et al., 2018; Honerkamp-Smith et al., 2009; Veatch et al., 2008; Veatch et al., 2007). If an IMP anchor has a marked affinity for the lipid phase constituting these fluctuating domains, it stabilizes in its neighbourhood such a nanodomain for energetic reasons, acting as a condensation nucleus, and giving rise to a long-lived lipid annulus around it. Contrary to the classical lipid-raft scenario where lipid nanodomains are relatively stable over time and exist independently of the presence of proteins, the lipid annuli can exist even though the surrounding lipids do not phase-separate spontaneously in the bulk, i.e. close to a critical point, however in the disordered phase. Given that the cell membrane is a multi-component lipid mixture, it is reasonable to speculate that several second-order demixing transitions occur simultaneously, giving rise to annuli of variable composition around different IMP anchors. If two IMP anchors localize in “like” and/or miscible annuli, they will tend to encounter with a higher probability because this condition reduces the interfacial energy cost at the annuli boundary (Gil et al., 1997; Katira et al., 2016). In contrast, if they localize in “unlike” and immiscible annuli, their close encounter will be less probable (i.e., less frequent), giving rise to a potentially rich combinatorial of protein/lipid interactions (Hinderliter et al., 2004). Here, we emphasize that the proximity of a second-order phase transition is not a very strong constraint in practice, since the correlation length for lipid mixtures of isolated cells membranes remains significantly larger than 1 nm even at temperatures of a few degrees above than the critical transition temperature (Veatch et al., 2008).

According to the simple sameness (“like”) rule, effective Lck trans-autophosphorylation should occur with a wide variety of anchor appended to Lck. The rule for distancing from CD45 should be modulated by the structural variance between Lck and CD45 anchors, but abolished when the two anchors are the same. More generally, our data show that the nature of the hydrophobic moieties embedding Lck and CD45 in the lipid bilayer influences the rate at which they interact laterally and of the underlying chemical reactions. Future studies should define if and to what degree the structural features of IMPs hydrophobic moieties modulate the balance between lateral proximity and distancing.

Crystallography and mass spectrometry have shown that membrane lipids can selectively bind to IMPs’ hydrophobic portions, with contributions of electrostatics or other polar interactions with TMD juxtaposed segments and/or ecto and cytosol moieties (Gupta et al., 2017; Laganowsky et al., 2014; Lee, 2011; Sun et al., 2018; Yen et al., 2018). Referred to as non-annular, these lipids can have structural roles (i.e., oligomerisation; allosteric regulation) and constitute also part of a first lipid layer, analogous to the protein hydration shell (Laage et al., 2017). Shape irregularities, polar amino acids and dimensions of MPs’ hydrophobic moiety cause considerable lipid bilayer deformation around the IMP to minimise free energy of solvation (Marsh, 2008; Mouritsen, 2010; Phillips et al., 2009). The energetically best-fitting lipids can be predicted to form the first shell(s) around the protein, due either to chemical affinity and/or pure physical criteria (e.g., lipid chain length/shape/unsaturation fitted to protein hydrophobic core). Once (a) first shell(s) is/are stabilized, it/they can recruit in turn a few more lipids because the involved lipid species tend to phase separate from the bulk. However, while annular lipids do not need to be associated with a complete phase-separated mixture, such a favourable environment close to the protein can promote a lipid condensation nucleus introduced above. From a molecular perspective, the result is a multilayer sheath that exhibits spatial and dynamic distribution distinct from bulk-solvent around the IMP (Marsh, 2008; Phillips et al., 2009).

The features of these local lipid have been captured by recent MDS of simple or very complex lipid mixtures embedding IMPs of known structure (Corradi et al., 2018; Corradi et al., 2019; Ingolfsson et al., 2022), leading to propose the notion of IMPs’ “lipid fingerprints” (Corradi et al., 2018), defining IMPs’ immediate lipid environment. Recent evidence using quasi-elastic neutron scattering has suggested that the helical TMD of the transferring receptor considerably reduces the dynamics of the surrounding lipids up to 5-10 nm distance, implying co-diffusion with the TMD, corroborated by MDS (Ebersberger et al., 2020), in full agreement with earlier MDS studies (Niemela et al., 2010). Recent progress in MS-based lipidomics of IMPs extracted in SMA-based native nanodisks (Teo et al., 2019) may be a promising avenue for experimentally define lipid fingerprints. The structure and dynamics of a lipid fingerprint surrounding IMPs necessarily leads to an interaction energy between them, determined by the sign and value of lipid mixing free energy, and resulting from the competition between lipid-lipid affinities and mixing entropy (Destainville et al., 2016). The energies at play will be moderate in the vicinity of criticality (Destainville et al., 2018; Gil et al., 1997; Honerkamp-Smith et al., 2009), nonetheless they are sufficient to reduce, though not abolish IMPs close proximity for immiscible annuli (Fig. **6A**). Conversely, two IMPs exhibiting the same annular lipids (i.e., each and every IMP with respect to itself) should experience an attractive interaction resulting in a higher probability for dynamic proximity (Fig. **6A**). This general property could prime the formation of IMP homo-clusters whose half-life should depend also on potential protein-protein interactions when proteins arrive at contact. In this respect, our data do not question the existence of membrane rafts intended as lipid-protein ensembles (Lingwood and Simons, 2010) driven by annular lipid sameness, such as IMPs homo-clusters (Destainville et al., 2016; Gil et al., 1998; Hinderliter et al., 2004), as exemplified by dynamic syntaxin clustering (Saka et al., 2014) or by more dynamic smaller clusters of Lck (Baumgart et al., 2016) or GPI-anchored proteins (Sharma et al., 2004), not necessarily formed in L_o_ membranes domains (Sevcsik et al., 2015). Recent high powered computationally MDS suggest that Ras dimers can occur in absence of protein-protein interaction (Ingolfsson et al., 2022).

**Figure 6.**
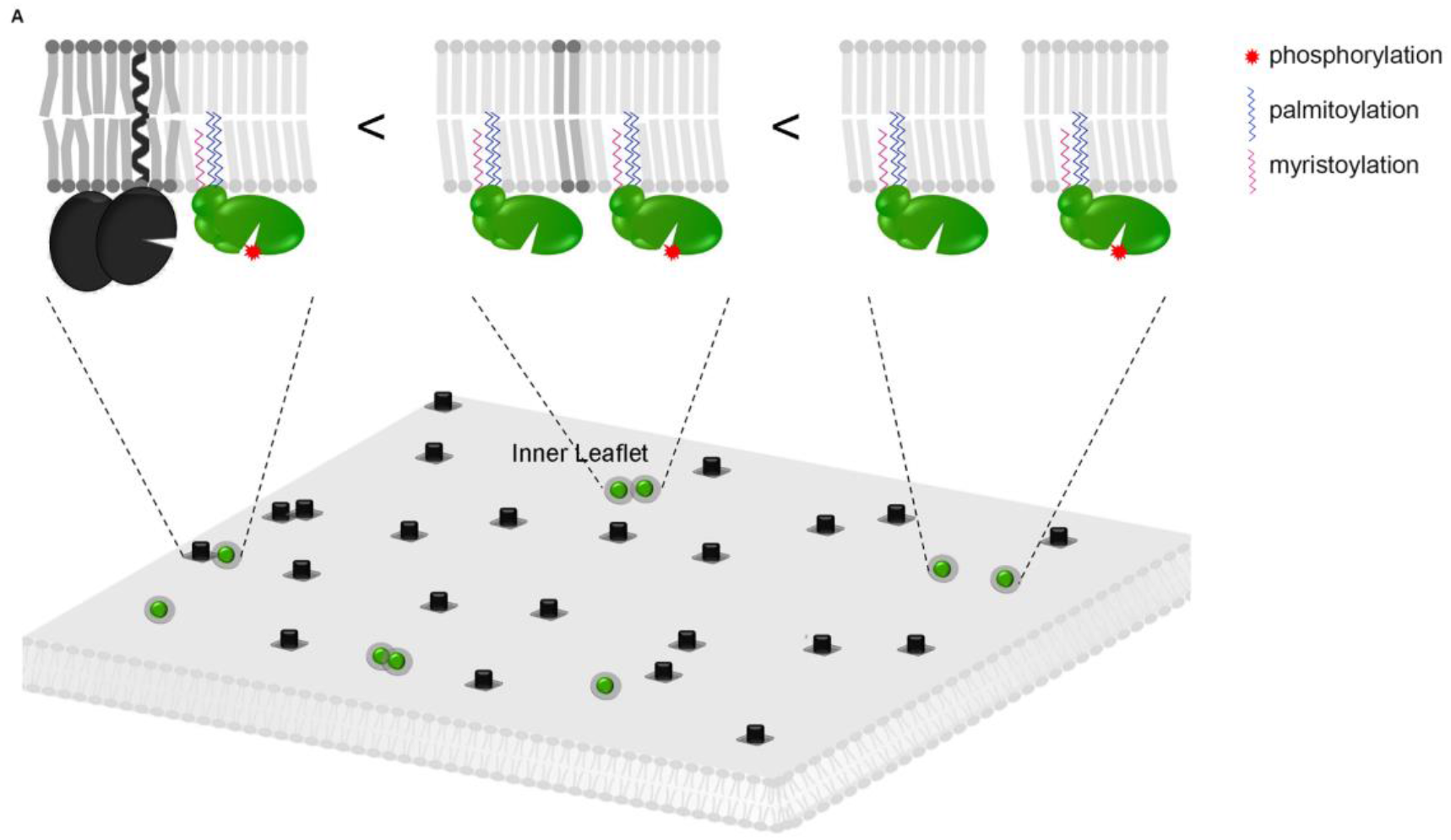
Schematic depiction of the “hydrophobic footprint” hypothesis. (**A**) The “hydrophobic footprint” is primarily made of annular lipids of specific composition and distribution co-diffusing with the membrane anchor. Different “hydrophobic footprints” create energetic barriers that reduce the probability of lateral proximity with potential functional consequences. (**Bottom**) “Like” annular lipids (light grey circle surrounding Lck - green) favour Lck:Lck pairing. “Unlike” annular lipids (dark grey squares surrounding CD45 - black) do not veto CD45:Lck pairing but make it less favourable. “Like” CD45:CD45 pairing may be inconsequential or have a regulatory role. (**Top**) The “hydrophobic footprint” for CD45 and Lck is idealised for simplicity by lipids of different aliphatic chain length and/or saturation, but can be further diversified by hydrophobic mismatch and/or interactions of IMP extracellular and cytoplasmic domains with lipid heads. The difference in probability of “hydrophobic footprint”-dependent close proximity may explain Lck_A_ formation and steady upkeep. Magenta and blue lines represent myristoylation and palmitoylation respectively; the red star indicates tyrosine phosphorylation.

## Author contributions

<u>Conceptualisation</u> by K.N. and O.A. <u>Project supervision</u> by K.N. and O.A. Most experiments were performed by N.P., D.C. and K.N., with additional ones by G.M. and A-L.L. <u>Supervision of SIM experiments, masks’ design and script for PM/CP</u> by L.S. <u>Supervision of ImageStream experiments, PM/CP script and data analysis</u> by D.H. and S.P.C. <u>Lck enzymatic activity modelling</u> by E.M., A.G. and M.D. <u>Data Interpretation and conceptual elaboration</u> by N.P., H-T.H., N.D., K.N. and O.A. <u>Manuscript writing</u>: original drafts by O.A., edited by N.P., K.N., N.D. and O.A. and the complete manuscript read by all authors.

## Acknowledgements

Wellcome Trust Grants GR076558MA and WT094296MA to O.A.; K. Karatheodori Program E609 (University of Patras) to K.N. Wellcome Trust Awards 091911 and 107457 to Micron Oxford; Ministero dell’Istruzione, Università e Ricerca (R. Levi-Montalcini fellowship) to M.D. Grant ANR-10-BLAN-1509 to H.-T.H. We thank Drs. Anna K. Schulze and Thomas Höfer (DKFZ, Heidelberg) for help with initial FCM data analysis and Prof. Simon Davis (Oxford University) for donation of a murine CD4 cDNA. We are particularly indebted with Drs. Antreas Kalli; Gerhard Schütz; Kai Simons, Ilpo Vattulainen, Rajat Varma, Peter Tieleman; Omer Dushek, Michael Dustin and Andres Alcover for helpful discussions and suggestions. We thank Christine Ralf and Ana Maria Vallés for reading the manuscript.

## METHODS

### Antibodies and reagents

Rabbit anti-pY505-Lck, anti-pY416-Src, rabbit anti-Lck (NBP1-85804) was from Novus Biologicals; anti-Lck-PE (73A5) antibodies (Abs) were from Cell Signaling Technology; rat anti-CD45 (YAML 501.4) Ab was from Santa Cruz Biotechnology; mouse anti-CD45-AF647 (HI30) Ab was from BioLegend. Secondary Abs for FACS analysis and SIM were: AlexaFluor 594 donkey anti-rat IgG, AlexaFluor 488 goat anti-rabbit IgG and AlexaFluor 647 goat antirabbit IgG (Thermo Fischer). A770041 was from Axon Medichem, Sodium Orthovanadate (Vanadate) was from New England BioLabs (NEB), catalase and hydrogen peroxide (30 %) from Sigma-Aldrich.

### Specificity of antibodies used for flow cytometry analysis and 3D-SIM

The specificity of the anti-pY416, anti-pY505 Abs has been extensively tested previously for immoboblot and for tissue staining (Nika et al., 2010). Here, we analysed further the reliability of the aforementioned Abs and of the anti-Lck 73A5 for flow cytometry and 3D-SIM. Induced or non-induced JCaM.1 cells expressing Lck were stained with anti-Lck 73A5-PE and anti-pY416 or anti-pY505 Abs, followed by anti-rabbit Alexa Fluor 647 and analysed by flow cytometry or 3D-SIM. Overlay histograms and 3D-SIM imaging in Fig. **S1C** and **S1F** show that all three antibodies exclusively reacted with dox-treated cells, which specifically express the Lck protein. Furthermore, reactivity of the anti-pY416 antibody, specifically recognizing pY394 of Lck (Nika et al., 2010) was lost after treatment of the induced cells with 2 μM A770041 or when the Ab was previously incubated with a synthetic peptide containing phospho-Y394 (Fig. **S1D**).

### IC_50_ evaluation of A770041

For Lck’ inhibition, we used A770041, which has a much greater specificity for Lck than for Csk (Burchat et al., 2006) but the IC_50_ for the latter is unknown. The IC_50_ of A770041 for Lck and Csk was determined by incubating serial dilution of the inhibitor with 1 μM of either recombinant Lck or Csk, 1 μM ATP and 1 μM substrate, as previously reported (Bain et al., 2007). A similar test was carried out using recombinant ZAP-70. Data shown in Table **S1** were obtained from MRC PPU Reagents and Services, School of Life Sciences (University of Dundee).

### Pervanadate preparation

The pervanadate (PV) solution was freshly prepared prior to each experiment as previously described (Huyer et al., 1997). Briefly, PV stock solution (1 mM) was prepared by adding 10 μl of 100 mM vanadate (NEB) and 50 μl of 100 mM hydrogen peroxide (diluted from a 30 % stock in 20 mM HEPES, pH 7.3) to 940 μl of H_2_O. Excess of hydrogen peroxide was removed by adding 200 μg/ml of catalase for 5 min after mixing the vanadate to the hydrogen peroxide. The PV solution was immediately used to minimize decomposition of the vanadate-hydrogen peroxide complex.

### Cells

Cell lines were maintained at 37 °C with 5 % CO_2_ in a humidified incubator (Heraeus). Human embryonic kidney epithelial Lenti-X293T (Clontech) cells were cultured in complete DMEM (Sigma Aldrich) supplemented with 15 % foetal bovine serum (FBS). Jurkat Cln20, JCaM.1 (both CD4- and CD8-negative) and JCaM.1-derived cell lines were cultured in RPMI 1640 supplemented with 10 % FBS. JCaM.1-derived cell lines with tetracycline-inducible gene expression system were maintained in RPMI 1640 supplemented with 10 % tetracycline-negative FBS (Clontech). Cells were routinely tested and found negative for mycoplasma and were not STR profiled.

Primary human CD4^+^ T cells (> 95 % pure) were isolated by negative selection from whole blood of healthy donors (National Blood Service, Bristol, UK) using the Dynal CD4 negative isolation kit (Thermo Fisher). Cells were routinely maintained in culture overnight in RPMI-1640, 10 % FBS before each experiment. For Lck inhibition, cells were treated with 2 or 5 μM A770041 (Axon) at 37 °C for 30 sec, 1 min or 5 min, as specified in the corresponding figure legend. For protein tyrosine phosphatase (PTP) inhibition, cells were treated with 100 μM Pervanadate (PV) for 1 or 3 min, as specified in the corresponding figure legend.

### Construction of chimeric or mutated proteins and cloning

The cDNA of human Lck wild-type (Lck) was used to generate all Lck chimeras and the cytoplasm-resident mutant LckΔSH4. All Lck constructs were cloned in the expression vector pLVX-Tight-Puro (Clontech Laboratories, Inc.), between 5’ NotI and 3’ EcoRI restriction sites. The SrcSH4-Lck chimera was generated by PCR using an oligonucleotide juxtaposing human SrcSH4 to human Lck. Specifically, the oligonucleotide used comprised the nucleotide sequence encoding amino acids 1-11 of human SrcWT, followed by amino acids 11-18 of Lck (Table **S2**). LckΔSH4 was obtained by PCR using a 5’ primer corresponding to amino acids 11-19 of Lck. To facilitate the generation of the LAT-, CD4-, CD4C/S- and CD45-Lck chimeric proteins, an XbaI restriction site was introduced prior to triplet coding for Asp11 of Lck. Then, NotI-XbaI fragments comprising the nucleotide sequences coding for the selected anchors were ligated to Lck XbaI-EcoRI fragment, lacking the SH4 domain (coding for residues 11-509) (see Table **S2**). The chimeras LAT-Lck and CD45-Lck were generated with cDNA of human LAT and human CD45 of our laboratories. For the CD4-Lck chimera we used as a template a cDNA of murine CD4 graciously provided by Prof Simon Davis laboratory. The CD4C/S-Lck chimera was generated in our laboratory by site-directed mutagenesis of our CD4-Lck construct. All chimeric and mutant constructs were verified by DNA sequencing.

### Production of lentiviral particles

Lentiviruses were generated using the packaging cell lines Lenti-X293T. The culture medium was exchanged with RPMI supplemented with 10 % FBS just prior to transfection. Lenti-X293T at 80 % confluence were transfected using PEIpro (Polyplus) according to the manufacturer’s instructions. The packaging plasmids pVSVG and pSPAX2 were mixed with the lentivirus expression vectors containing the gene of interest. PEIpro solution was added to the plasmids mix and immediately vortexed, left 15 min at room temperature (RT) and then added dropwise to the cells by gently swirling the plate. Supernatant containing lentiviral particles was collected after 48 h and filtered through a 0.45 μm sterile filter (Sartorius Stedim). Lentivirus supernatants were concentrated with PEG-*it*™ (SBI) concentration kit according to the manufacturer’s instruction. Briefly, lentiviral supernatants were mixed with Virus Precipitation Solution (SBI) to a final concentration of 1X Virus Precipitation Solution and incubated overnight at 4 °C followed by a centrifugation at 1,500 x g for 30 min at 4 °C. Pellets containing lentivirus particles were re-suspended in 1/100 of the volume of the original cell culture using cold RPMI. Aliquots were immediately frozen in cryogenic vials at - 80 °C and stored until use. Aliquots of each lentivirus batch were routinely pre-tested by serial dilution titration. Frozen aliquots were thawed only once and used immediately with minimal loss of virus titre as determined by flow cytometry.

### Generation of Tet-On inducible cell lines

Stable, inducible cell lines were generated using the Lenti-X Tet-On-Advanced Inducible Expression System (Clontech Laboratories, Inc.) according to the manufacturer’s instructions. Briefly, JCaM.1 were transduced with lentiviral particles (as described above) containing the PLVX-Tet-On-Advanced vector, which constitutively expresses the tetracycline-controlled trans-activator rtTA-Advanced. 48 h after transduction, the transduced cells were put under selection by Geneticin (1 mg/ml) to generate a stable JCaM1-TetON cell line. This parental cell line was then transduced with lentiviral particles of pLVX-Tight-Puro containing the Lck constructs and, 48 h after transduction, put under selection by Puromycin (10 μg/ml) and Geneticin (1 mg/ml) to generate the respective stable cell line. Expression of the Lck constructs was induced by 1 μg/ml doxycycline (dox, Sigma-Aldrich) added to the cell culture medium, routinely 16 - 18 h prior to each experiment (Over Night (ON) dox-inductions).

### CellTrace violet labelling

To quantitatively evaluate the formation of Lck_A_ depending on Lck_T_ and according to different lipid anchor, we employed a flow cytometry-based approach that allows to concomitantly detect Lck_A_ and Lck_T_ on a per-cell basis. To improve precision and accuracy, we performed the double staining of Lck_A_ and Lck_T_ of two different JCaM.1-derived cell lines expressing mutated or chimeric-Lck together with the JCaM1-Lck cell line (used as an internal reference). To this aim, we labelled the two cell lines with different concentration of CellTrace violet (1 and 0.25 μM, Thermo Fisher) and the JCaM1-Lck cell line with a carrier control (DMSO, Sigma) prior the ON dox-induction. Specifically, cells were washed once in PBS and adjusted to a final concentration of 10^6^ cells/ml in pre-warmed PBS at 37 °C. CellTrace violet (Thermo Fisher) or carrier control DMSO (Sigma) was added to reach the staining concentration indicated above and cells were incubated at 37 °C in the dark. After 20 min, samples were diluted 5-fold in complete medium and incubated for an additional 5 min at 37 °C in the dark. After removal of the diluted staining solution, cells were re-suspended in complete medium, counted, mixed and induced, in the same well, for Lck (WT, mutated or chimeric-Lck) by ON-addition of 1 μg/ml dox.

### Flow cytometry

For cell surface staining, single-cell suspensions were transferred into a 96-well V-bottom plate, washed once with 100 μl FACS buffer (0.5 % bovine serum albumin (BSA) (Sigma) in PBS). After spinning, supernatants were removed and cell pellets re-suspended in 50 μl staining solution containing fluorescence-conjugated primary Ab diluted in FACS buffer and incubated for 20 min at RT. Cells were then washed twice and either acquired immediately on a FACS Calibur flow cytometer (BD Biosciences) or BD LSR Fortessa X20 (BD Biosciences) or fixed with a pre-warmed fixation solution (BD Cytofix®, BD Biosciences) for 10 min at 37 °C. Cells were then washed twice in 150 μl permeabilisation buffer (BD Perm/Wash I, BD Biosciences), re-suspended in 150 μl permeabilisation buffer and incubated at 4 °C for 30 min. Primary antibodies, diluted in permeabilisation buffer, were added to the cells for 1 h, followed by three washes in permeabilisation buffer and the addition of the corresponding secondary antibodies (in permeabilisation buffer). After 3 washes, cells were analysed on a FACS Calibur flow cytometer (BD Biosciences) or BD LSR Fortessa X20 (BD Biosciences). Acquired data were analysed by FlowJo (FlowJo Software part of BD). Counts, percentages or median intensity fluorescence values (MFI) were extracted from FlowJo as excel files. Statistical analysis and non-linear regression were performed with Prism (GraphPad Software).

### Imaging Flow Cytometry (ImageStream)

Samples were stained for Lck, CD45 and DAPI according to the general protocol for intracellular staining described above (see Flow Cytometry section). After staining, cells were re-suspended at 1*10^7^ cells per ml for loading onto the ImageStream instrument. Samples were run on a 2 camera, 12 channel ImageStream X MkII (Amnis Corporation) with the 60X Multimag objective, the extended depth of field (EDF) option providing a resolution of 0.3 μm per pixel and 16 μm depth of field. Bright field images were captured on channels 1 and 9 (automatic power setting). At least 10,000 images per sample were acquired using INSPIRE 200 software (Amnis Corporation) and then analysed using the IDEAS v 6.2 software (Amnis Corporation). A colour compensation matrix was generated for all the fluorescence channels using samples stained with single colour reagents or antibody conjugate coated compensation beads, run with the INSPIRE compensation settings, and analysed with the IDEAS compensation wizard. Images were gated for focus (using the Gradient RMS feature) on both bright field channels (1 and 9) followed by selecting for singlet cells (DNA intensity/aspect ratio). A mask depicting the plasma membrane (PM) was defined from the anti-CD45 staining, used as a membrane marker, and a ratio between the MedianFI of Lck at the PM and the MedianFI of Lck in the rest of cell was calculated.

### Immunostaining, 3D-SIM, image acquisition and analysis

Single-cell suspensions were immobilized on poly-L-lysine (Sigma-Aldrich)-coated high No. 1.5H precision glass coverslips (Marienfeld-Superior) for 10 min at 37 °C in a cell culture incubator. Cells were fixed for 10 min with 4 % formaldehyde/PBS and permeabilised with 0.5 % Triton-X100/ PBS for 5 min. After blocking with PBS/1 % BSA for 15 min, cells were stained for 1 h at room temperature (RT) with the indicated primary antibodies. Fluorochrome-conjugated secondary antibodies were added for 1 h. Nuclei were counterstained with 1 μg/ml DAPI (Sigma-Aldrich) and coverslips were mounted to microscopy slides with ProLong Gold anti-fade reagent (Thermo Fisher). 3D-SIM was performed on an OMX V3 Blaze microscope (GE Healthcare) using 405-, 488- and 592-nm laser lines and a 60x/1.42 oil UPlanSApo objective (Olympus). Multi-channel images were captured sequentially by sCMOS cameras (PCO). 1 μm stacks were acquired at 125 nm z-distance, with 15 raw images per plane (three angles, five phases) resulting in 120 raw images in total, for each sample. Calibration measurements of 0.2 μm diameter TetraSpeck fluorescent beads (Thermo Fisher) were used to obtain alignment parameters subsequently utilized to align images from the different colour channels. Image stacks were computationally reconstructed from the raw data using the SoftWoRx 6.0 software package (GE Healthcare) to obtain super-resolution image with a resolution of wavelength-dependent 100-130 nm in x and y and 300-350 nm in z. Raw and reconstructed image data quality was confirmed using *SIMcheck* ImageJ plugin (Ball et al., 2015). Image processing and evaluation was performed using in-house ImageJ scripts: 32-bit reconstructed image stacks were thresholded to the modal intensity value (defining the centre of noise) and converted to 16-bit composites. The central four image planes were then average projected and Gaussian blurred (sigma 3 pixel). Regions of interest (ROI) covering the nuclear and plasma membrane (PM) were defined by “Otsu” auto-thresholding in the DAPI and anti-CD45 channel, respectively, and applying further processing steps (“Binary mask”, “Fill holes” and “Erode”). The area between the PM and nuclear ROI was defined as the cytoplasmic ROI. Measurements of the average fluorescence intensity within the respective PM and cytoplasm ROIs were used to calculate the PM/cytoplasm ratios for the staining of anti-Lck, anti-Src, anti-pY416 and anti-pY505 antibodies.

### FACS analysis of Lck_A_ generation as a function of Lck_T_

Cell expressing Lck, Lck-chimeras or mutant were concomitantly stained for Lck_A_ and Lck_T_ as described above (see Flow Cytometry section). The corresponding flow cytometry 2D plots were linearised by converting the *x* (Lck_T_) and *y* (Lck_A_) axes from a logarithmic to a linear scale to increase resolution and a dense binning (n > 33) was applied to the entire population, within a physiological concentration range of Lck_T_ set on Cln20. Values of the geometric median and its associated SD for Lck_A_ and Lck_T_ in each bin, were extracted. Background-subtracted values of the geometric median for Lck_A_ and Lck_T_ in each bin were subjected to regression analysis.

### Probabilistic Model of Lck_A_ formation

To investigate Lck_A_ formation as a function of Lck_T_, we generated a simple probabilistic model where Lck can assume three different states: the inactive conformation (Lck_I_), the primed conformation (Lck_P_) and the active conformation (Lck_A_). Therefore, we consider the following reactions:

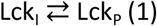

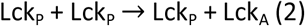

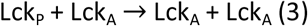

The assumptions used in the model are as following:

- In the initial state, the first equilibrium reaction is shifted towards the primed conformation.
- We consider two different probabilities for (2) and (3).
- The increase of total Lck (Lck_T_) is included in the model by the presence of an additional parameter.
- The effect of CD45 is not included in the model.

Starting from these considerations, we studied the variation of Lck_A_ depending on Lck_T_. For each cycle, each Lck can interact with any other Lck and this interaction can either i) keep the situation unchanged (Lck_P_ + Lck_P_ → Lck_P_ + Lck_P_) or ii) activate one of the two molecules (Lck_P_ + Lck_P_ → Lck_P_ + Lck_A_). Subsequently, another reaction takes place: as the presence of Lck_A_ increases, Lck_A_ and Lck_P_ can interact with each other to generate two active forms. The probabilities associated to these reactions (1) and (2), P_PA_ and P_AA_ respectively, are variable during the simulations (from 0.1 to 1 with steps of 0.05).

---

**Table S1.**
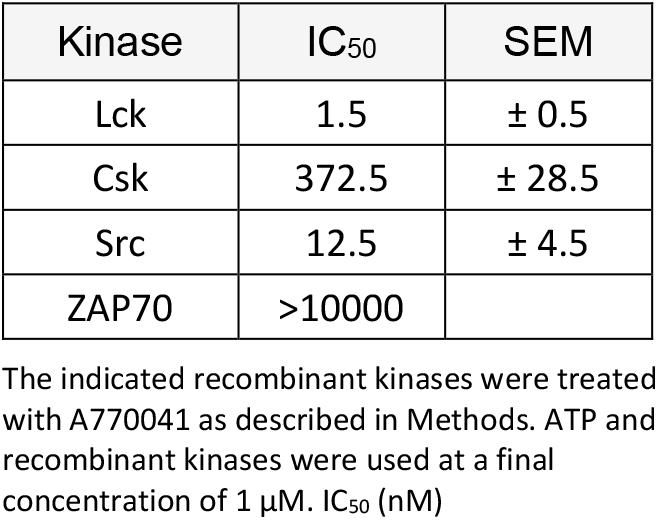
A770041 IC_50_.

**Table S2.** Amino acid sequence of Lck chimera’s membrane anchors.

| Chimeric proteins | AA sequence fused to LckΔSH4<sup>1</sup> | Lateral diffusion<sup>2</sup> | References |
| --- | --- | --- | --- |
| SrcSH4-Lck | **GSNKS**KPKDA | R/SRE | Smith et al., 2016 |
| LAT-Lck | EEA**ILVPCVLG**LLLL**PILAMLMALCVHCHRLPGS** | R/SRE | Douglass and Vale, 2005 |
| CD4-Lck | EEA**VFLACVLGGSFGFLGFLGLC****ILCCVRCR** | R/SRE | (Foti et al., 2002; Grebenkamper and Nicolau, 1995) |
| CD4C/S-Lck | EEA**VFLACVLGGSFGFLGFLGLC****ILCSVRSR** | Unknown |  |
| CD45-Lck | EEA**ALIAFLAFLIIVTSIALLVVLY** | R/HD | (Cairo et al., 2010) |

<sup>1</sup> All chimeras lack the sequence GCGCSSHPED, corresponding to LckSH4. Bold characters: membrane anchor chosen for each chimera with the putative TM region (underlined) according to UniProtKB. Each membrane anchor was fused to LckΔSH4, whose N-terminal sequence begins as DWMENIDVC. EEA is the short extracellular sequence of LAT (regular characters) introduced to facilitate similar chimera expression.
<sup>2</sup> Lateral diffusion behaviour of the anchor-donor proteins as reported in the indicated references.
R: random; SRE: Short-range entrapment; HD: hop diffusion. All the sequences are human, except the CD4 anchor region that is from the murine protein.

