## Supplementary Figures for "A role of Lck annular lipids in the steady upkeep of active Lck in T cells"

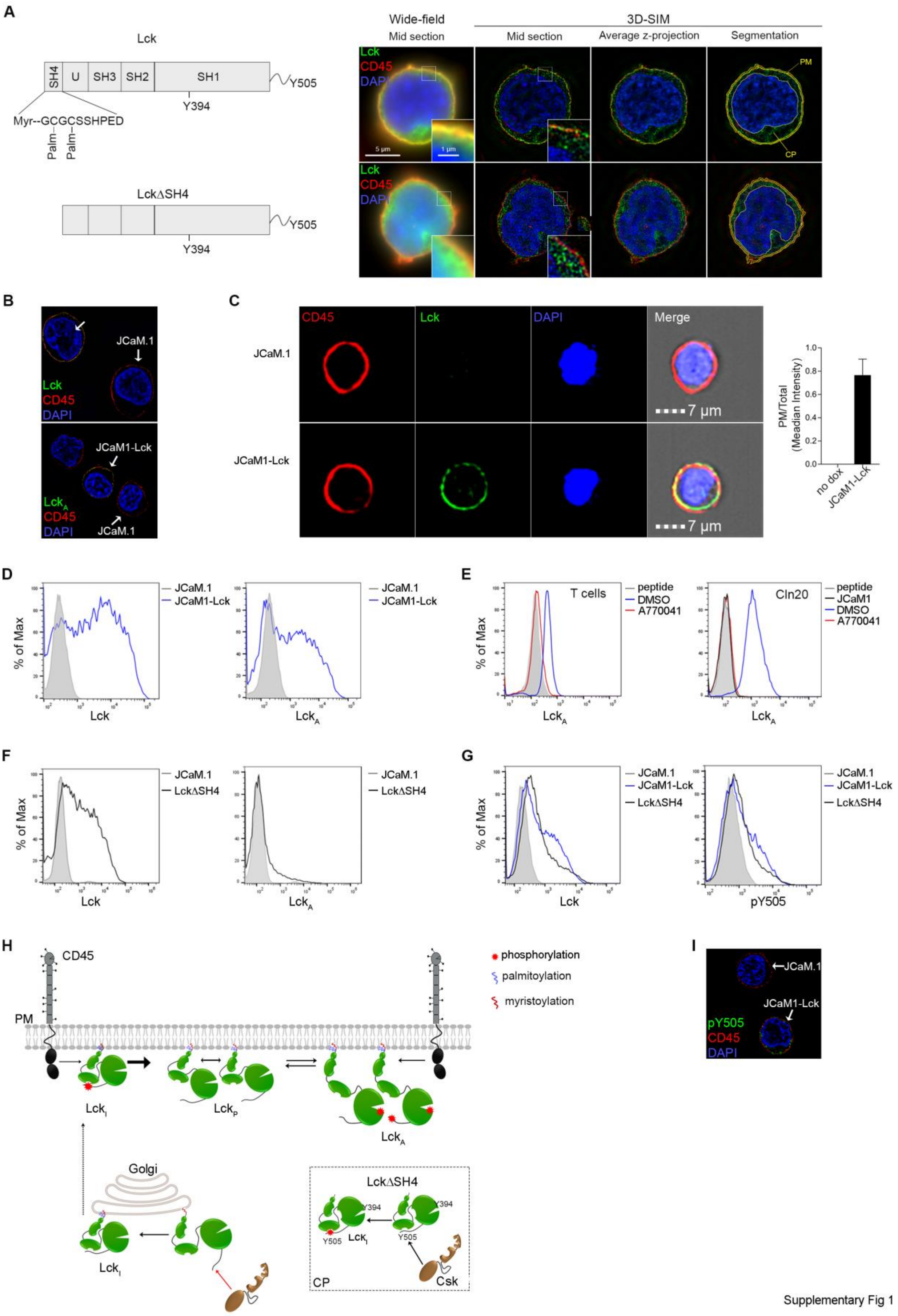


Supplementary Fig 1

**Figure S1. Related to Fig. 1**

(**A**) **Left**, schematic representation of Lck (**top**) and Lck∆SH4 (**bottom**) employed in this investigation. **Right**, Comparison of wide-field, 3D-SIM imaging and principles of quantitative image analysis for Lck (**top**) and Lck∆SH4 (**bottom**) subcellular localisation. Plasma membrane (PM) and Cytoplasmic (CP) region of interest are segmented in the central region of 3D image stack and average fluorescence intensity determined. Lck (green), CD45 (red) and DAPI (blue). CD45 and DAPI are used as a membrane and nuclear marker respectively. (**B**) 3D-SIM imaging showing the fidelity of the anti-pY416 and the anti-Lck antibodies (Abs) employed to specifically detect $Lck_A$ and Lck respectively. **Top**, 3D-SIM imaging of Lck (green) and **bottom**, $Lck_A$ (green) in a field containing a mix 1:1 of both non-dox-induced JCaM.1 and dox-induced JCaM.1 expressing Lck cells. Membrane marker CD45 (red), Nuclear marker DAPI (blue). (**C**), ImageStream of Lck (green) in JCaM.1 (negative control) or JCaM.1 expressing Lck. Scale bar: 7 µm. Membrane marker CD45 (red), Nuclear marker DAPI (blue). **Right**, quantitative analysis of Lck distribution at the Plasma Membrane (PM). Bars represent PM/Total ratio of Lck (Median). Error bars: MAD for n ≥ 5000 cells from 3 independent experiments. (**D**) Flow CytoMetry (FCM) showing the fidelity of the anti-pY416 and the anti-Lck antibodies (Abs) employed to specifically detect $Lck_A$ and Lck respectively by FACS analysis. Representative FCM histograms of Lck (**left**) and $Lck_A$ (**right**) in non-dox (doxycycline)-induced JCaM.1 (grey, line-filled area) and dox-induced JCaM.1 expressing Lck (blue). (**E**) FCM plots of $CD4^+$ T cells (**left**) and Cln20 (**bottom**) showing the fidelity of the anti-pY416 Ab when competing for anti-pY416 binding with a pY394-containing synthetic peptide. **Left**, representative FCM histogram of $Lck_A$ in $CD4^+$ T cells treated with 5 µM A770041 (red) or carrier (DMSO, blue) for 5' at 37 °C and stained with anti-pY416 or with anti-pY416 previously incubated with a pY394-containing synthetic peptide (grey, line-filled area). **Right**, representative FCM histogram of $Lck_A$ in Cln20 treated with 5 µM A770041 (red) or carrier (DMSO, blue) for 5' at 37 °C and stained with anti-pY416 or with anti-pY416 previously incubated with a pY394-containing synthetic peptide (grey, line-filled area). JCaM.1 were used as a negative control to set pY416 Ab background. (**F**) Representative FCM histograms of Lck (**left**) and $Lck_A$ (**right**) in non-dox-induced JCaM.1 (grey, line-filled area) and dox-induced JCaM.1 expressing Lck∆SH4 (black). Non-dox-induced JCaM.1 were used as a negative control to set Abs background. (**G**) Representative FCM histograms of Lck (**left**) or pY505-Lck (**right**) in JCaM.1 expressing Lck (blue) or LckΔSH4 (black). JCaM.1 (grey, line-filled area) were used as a negative control to set Abs background. (**H**) Schematic representation of Lck regulation at either the plasma membrane (PM) or the cytoplasm (CP). In the CP, Lck is synthetised and converted in $Lck_I$ (inactive) by Csk that phosphorylates Y505-Lck presumably in an exocytic compartment *en route* to the PM. Once at the PM, $Lck_A$ is generated and maintained by the solely antagonism between CD45 and Lck for Y394 phosphorylation. Specifically, at the PM, $Lck_I$ is rapidly dephosphorylated at Y505 by CD45 and converted in $Lck_P$. $Lck_P$ in turn generates $Lck_A$ that can either be dephosphorylated again by CD45 or accumulates at the PM. On the contrary, Csk controls $Lck_I$ almost exclusively in the CP as confirmed by LckΔSH4 mutant that, lacking the Lck lipid anchor, is only CP-resident. CP-resident LckΔSH4 is phosphorylated at Y505 by Csk, but being incapable of PM localization, cannot generate $Lck_A$. (**I**) 3D-SIM imaging of pY505-Lck (green) in a field containing a mix 1:1 of both non-dox-induced JCaM.1 and dox-induced JCaM.1 expressing Lck cells. Membrane marker CD45 (red), Nuclear marker DAPI (blue).

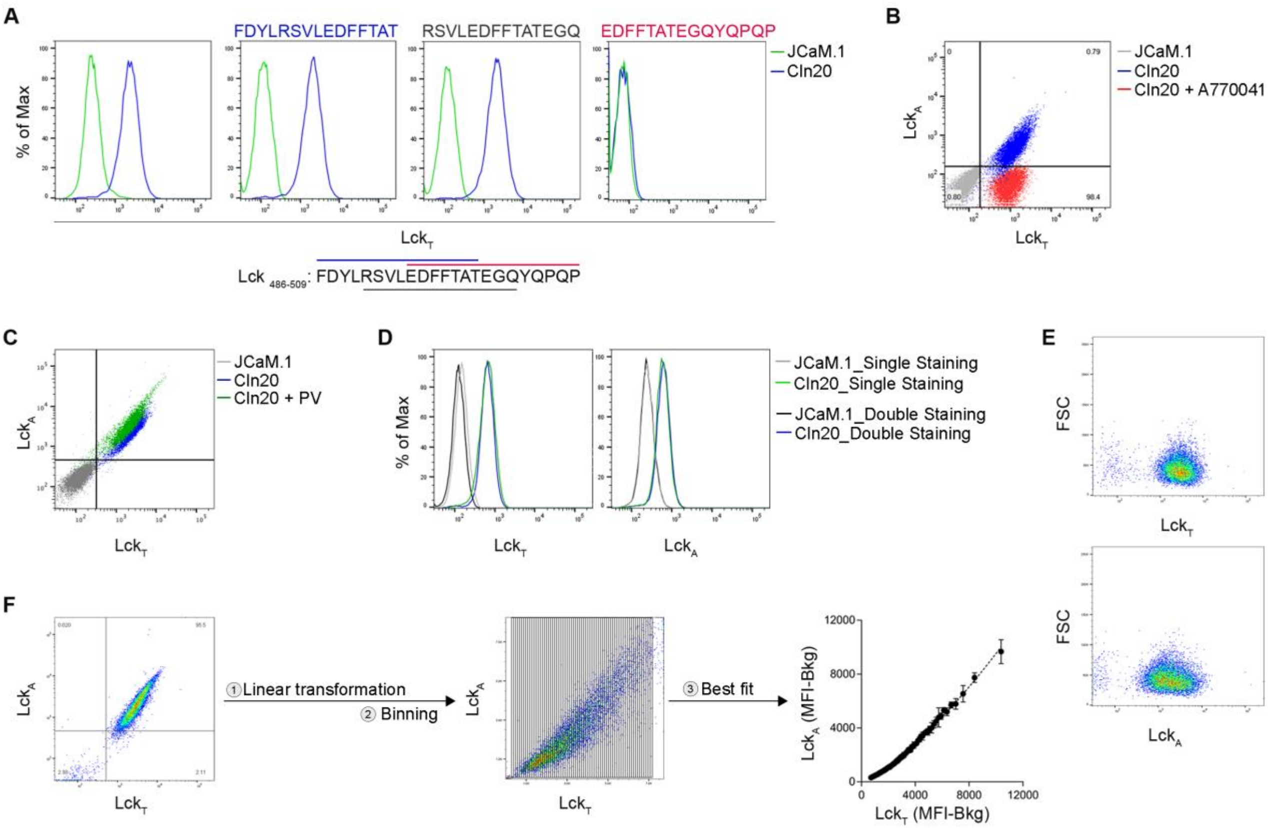


**Figure S2. Related to Fig. 2**

(**A**) Epitope mapping of the anti-Lck Ab (73A5) used to detect total Lck (Lck$_T$) in this investigation. **Top**, FCM histograms of Lck$_T$ in JCaM.1 (green) and Cln20 (blue) stained with 73A5 Ab pre-incubated or not with the competing peptides for the anti-Lck binding site whose aminoacidic sequence is indicated above each plot. JCaM.1 were used to assess Ab background. **Bottom**, sequence of Lck C-terminal tail used to raise the 73A5 Ab and showing the competing peptides' sequence and localisation. (**B**) Representative FCM 2D plot of Cln20 concomitantly stained for Lck$_A$ and Lck$_T$ in cells treated (red) or not (blue) with 2 $\mu$M of A770041 for 1 min at 37 $^{o}$C. JCaM.1 (grey) were used to assess Abs background. (**C**) Representative FCM 2D plot of Cln20 concomitantly stained for Lck$_A$ and Lck$_T$ in cells reacted (green) or not (blue) with catalase-treated pervanadate (PV) for 1' at 37 $^{o}$C. JCaM.1 (grey) were used to assess Abs background. (**D**) **Left**, representative FCM histogram of Lck$_T$ in Cln20 stained by anti-Lck 73A5 in presence (blue, double staining) or absence (green, single staining) of anti-pY416. JCaM.1 (grey and black) were used to assess Ab background. **Right**, representative FCM histogram of Lck$_A$ in Cln20 stained by anti-pY416 in presence (blue, double staining) or absence (green, single staining) of anti-Lck 73A5. JCaM.1 (grey and black) were used to assess Ab background. (**E**) **Top**, representative FCM 2D plot of $Lck_T$ depending on cell size (FSC, Forward SCatter). **Bottom**, representative FCM 2D plot of $Lck_A$ depending on cell size (FSC, Forward SCatter). (**F**) Flow chart depicting the experimental procedure followed to uncover $Lck_A$ dependence on $Lck_T$. **Left**, representative FCM 2D plot of Cln20 concomitantly stained for $Lck_A$ and $Lck_T$. **Middle**, after converting *x* ($Lck_T$) and *y* ($Lck_A$) axes from a logarithmic to a linear scale, a dense binning (n = 73) was applied to the entire population and the values of the geometric median for $Lck_A$ and $Lck_T$ in each bin, were extracted. **Right**, background-subtracted values of the geometric median for $Lck_A$ and $Lck_T$ in each bin were subjected to regression analysis. Non-linear regression fit of $Lck_A$ (MFI-Bkg) vs. $Lck_T$ (MFI-Bkg), n = 2, $R^2 = 0.99$; F-test $p < 0.0001$.

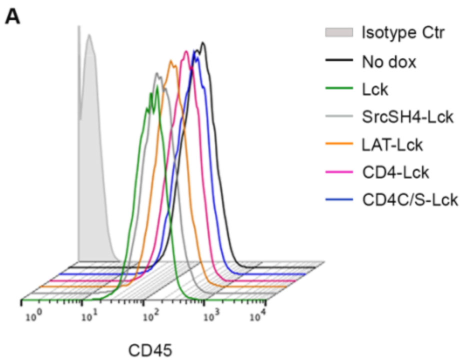


**Figure S3. Related to Fig. 3**

(**A**) CD45 detection in JCaM.1 expressing Lck or the indicated Lck chimera. Representative FCM histograms of CD45 by anti-CD45 Ab in non-dox-induced JCaM.1, JCaM.1 expressing Lck (green), Src-SH4 (grey), LAT-Lck (orange), CD4-Lck (magenta) or CD4C/S-Lck (blue). An isotype control was used to set the Ab background (grey, line-filled area).

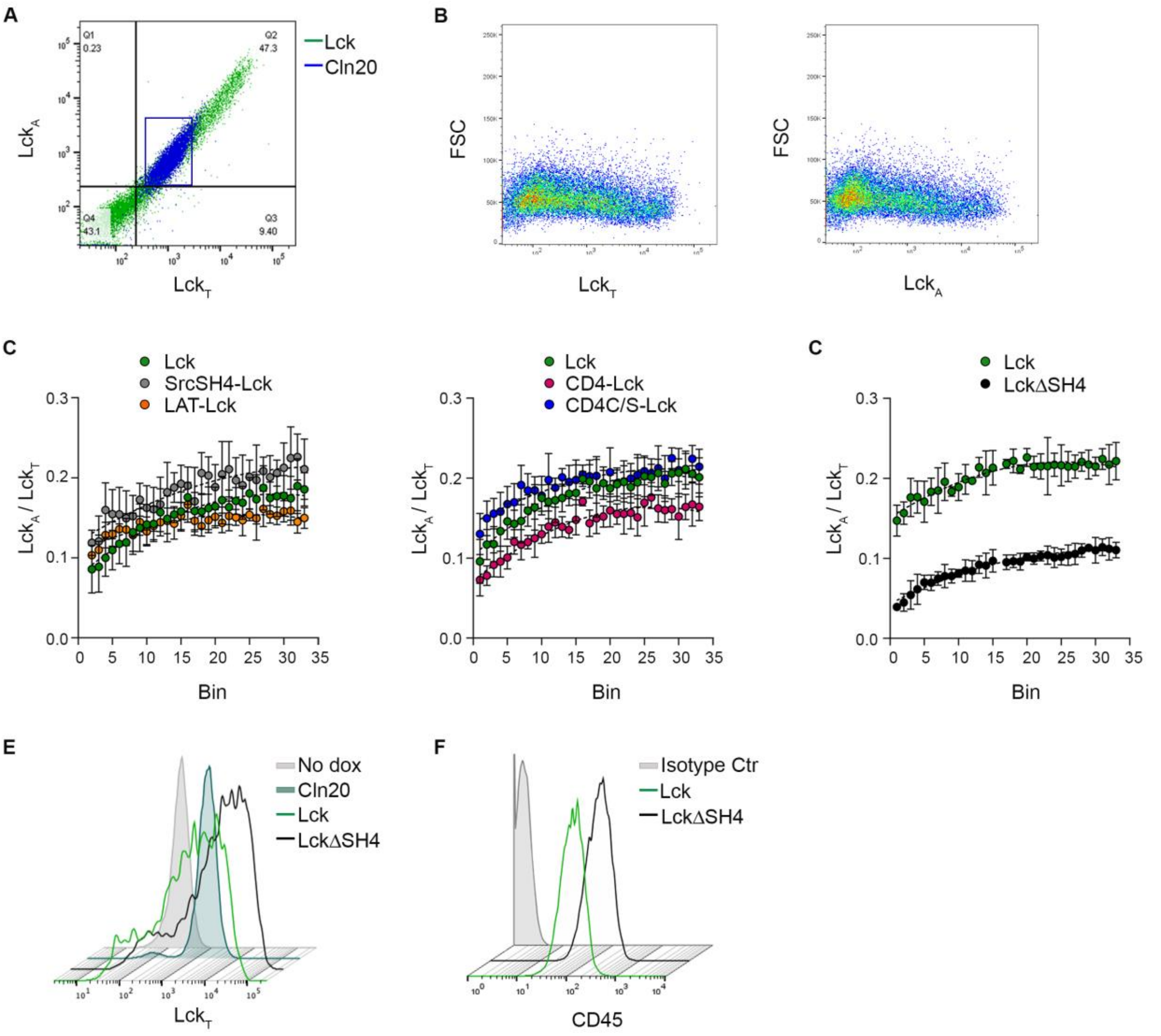


**Figure S4. Related to Fig. 4**

(**A**) Representative FCM 2D plot of JCaM.1 expressing Lck (green) and Cln20 (blue). A gate (blue box) containing the limits of Lck expression in Cln20 was applied to set a physiological concentration range of $Lck_T$ to analyse $Lck_A$ dependence on $Lck_T$. Cells were labelled (Cln20) or not (JCaM.1-Lck) with CellTrace violet, induced for Lck expression by dox and 16/18 h after, analysed by FACS. N = 3, staining in duplicate. (**B**) **Left**, representative FCM 2D plot of $Lck_T$ depending on cell size (FSC, Forward SCatter). **Right**, representative FCM 2D plot of $Lck_A$ depending on cell size (FSC, Forward SCatter). (**C**) $Lck_A$ formation depending on different membrane anchors, in JCaM.1 expressing Lck or the indicated Lck chimera or mutant. Cells were labelled or not with two different concentrations of CellTrace violet, mixed 1:1:1, induced for Lck expression by dox and, 16/18 h after, concomitantly analysed by FACS for $Lck_A$ and $Lck_T$. A dense binning (n = 33) within a physiological concentration range of $Lck_T$ set on Cln20, was applied and the values of the geometric median for $Lck_A$ and $Lck_T$ in each bin, were extracted (see **Fig. 4B**, left panels) and plotted as a ratio ($Lck_A$/ $Lck_T$). 2D plots showing $Lck_A$ formation normalised by $Lck_T$ ($Lck_A$/ $Lck_T$ ratio) in each bin in (**left**) JCaM.1 cells expressing Lck (green), SrcSH4-Lck (grey) or LAT-Lck (orange) and (**right**) Lck (green), CD4-Lck (magenta) or CD4C/S-Lck (blue). Each bin represents a sequential increase of $Lck_T$. N = 3, staining in duplicate. (**D**) $Lck_A$ formation depending on Lck membrane anchors. Cells were labelled or not with CellTrace violet, mixed 1:1, induced for Lck expression by dox and, 16/18 h after, concomitantly analysed by FACS for $Lck_A$ and $Lck_T$. A dense binning (n = 33) within a physiological concentration range of $Lck_T$ set on Cln20, was applied and the values of the geometric median for $Lck_A$ and $Lck_T$ in each bin, were extracted (see **Fig. 4C**, left panel) and plotted as a ratio ($Lck_A$/ $Lck_T$). 2D Plot showing $Lck_A$ formation normalised by $Lck_T$ ($Lck_A$/ $Lck_T$ ratio) in each bin in JCaM.1 cells expressing Lck (green) or LckΔSH4 (black). Each bin represents a sequential increase of $Lck_T$. N = 3, staining in duplicate. (**E**) Representative FCM histograms of $Lck_T$ by 73A5 staining of non-dox induced JCaM.1 (grey, line-filled area used as a negative control) or dox-induced JCaM.1 to express Lck (green) or LckΔSH4 (black) and Cln20 (dark green, line-filled area used to set a physiological range of $Lck_T$). (**F**) CD45 detection in JCaM.1 expressing Lck or LckΔSH4. Representative FCM histograms of CD45 by anti-CD45 Ab in JCaM.1 expressing Lck (green) or LckΔSH4 (black). An isotype control was used to set the Ab background (grey, line-filled area).

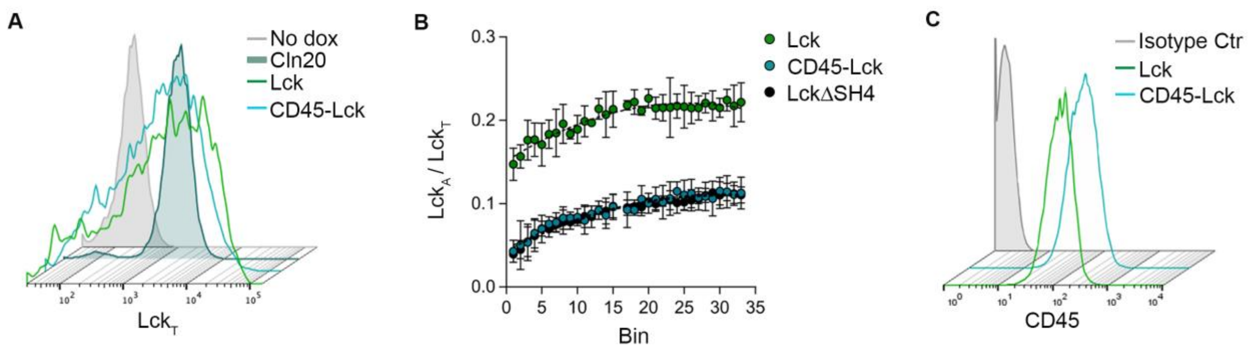


**Figure S5. Related to Fig. 5**

(**A**) Lck detection in JCaM.1 expressing Lck or CD45-Lck and Cln20. Cells were labelled or not with CellTrace violet, mixed and analysed by FACS for Lck expression. Representative FCM histograms of $Lck_T$ by 73A5 staining of non-dox induced JCaM.1 (grey, line-filled area, used as a negative control) or dox-induced JCaM.1 to express Lck (green) or CD45-Lck (cyan) and Cln20 (dark green, line-filled area used to set a physiological range of $Lck_T$). (**B**) $Lck_A$ formation depending on different membrane anchors, in JCaM.1 expressing Lck or the indicated Lck chimeras or mutant. Cells were labelled or not with two different concentrations of CellTrace violet, mixed 1:1:1, induced for Lck expression by dox and, 16/18 h after, concomitantly analysed by FACS for $Lck_A$ and $Lck_T$. A dense binning (n = 33) within a physiological concentration range of $Lck_T$ set on Cln20, was applied and the values of the geometric median for $Lck_A$ and $Lck_T$ in each bin, were extracted (see Fig. **5C**, left panel) and plotted as a ratio ($Lck_A$/ $Lck_T$). 2D Plot showing $Lck_A$ formation normalised by $Lck_T$ ($Lck_A$/ $Lck_T$ ratio) in each bin in JCaM.1 cells expressing Lck (green), CD45-Lck (cyan) or LckΔSH4 (black). Each bin represents a sequential increase of $Lck_T$. N = 3, staining in duplicate. (**C**) CD45 detection in JCaM.1 expressing Lck or CD45-Lck. Representative FCM histograms of CD45 by anti-CD45 Ab in JCaM.1 expressing Lck (green) or CD45-Lck (cyan). An isotype control was used to set the Ab background (grey, line-filled area).
